# Gas6 signaling via its cellular receptors is essential for homeostasis in response to endotoxemia and infection

**DOI:** 10.64898/2026.11.13.766121

**Authors:** Sara Çaku, Raja Prince, Laurent Burnier, Bojun Li, Thierry Roger, Marc Chanson, Rim Diab, Ruth Etzensperger, Mariantonietta Tripodo, Tanja Knopp, Elisa Introini, Lucas Liaudet, Joseena Mariam Iype, Charithani B. Keragala, Robert L. Medcalf, Carmen A. Pfortmueller, Ramanjaneyulu Allam, Anna S. Messmer, Anne Angelillo-Scherrer

**Affiliations:** Department of Hematology and Central Hematology Laboratory, Bern University Hospital, Inselspital, University of Bern, Bern, Switzerland; Department for BioMedical Research, University of Bern, Bern, Switzerland; Service and Central Laboratory of Hematology, Centre Hospitalier Universitaire Vaudois and University of Lausanne, Lausanne, Switzerland; Infectious Diseases Service, Centre Hospitalier Universitaire Vaudois and University of Lausanne, Lausanne, Switzerland; Department of Cell Physiology and Metabolism, Faculty of Medicine, University of Geneva, Geneva, Switzerland; Service of Adult Intensive Care Medicine, Centre Hospitalier Universitaire Vaudois and University of Lausanne, Lausanne, Switzerland; University Institute of Clinical Chemistry, Bern University Hospital, University of Bern, Bern, Switzerland; Molecular Neurotrauma and Haemostasis, Australian Centre for Blood Diseases, Monash University, Melbourne, Australia; Department of Intensive Care Medicine, Inselspital, Bern University Hospital, University of Bern, Bern, Switzerland

**Keywords:** Gas6, Tyro3, Axl, Mer, cytokines, endotoxemia, sepsis, innate immunity

## Abstract

Sepsis is a life-threatening syndrome characterized by infection-induced organ dysfunction and dysregulated host inflammatory responses. Despite advances in supportive care, effective pharmacological therapies remain lacking. Growth arrest-specific protein 6 (Gas6), a vitamin K-dependent ligand of the TAM receptors (Tyro3, Axl, and Mer), operates at the interface of inflammation and coagulation. Here, we investigated the role of Gas6–TAM signaling in endotoxemia and infection in humans and mice.

Plasma Gas6 levels increased rapidly during sepsis in patients, during experimental endotoxemia in both humans and mice, and during infection in mice, and these elevations correlated with disease severity and poor clinical outcomes. In both species, circulating Gas6 remained closely balanced with soluble Axl (sAxl), suggesting that much of plasma Gas6 is sequestered in soluble complexes, thereby limiting activation of membrane TAM receptors.

In mice, deficiency of Gas6 or individual TAM receptors increased susceptibility to endotoxemia, and Gas6 deficiency additionally heightened vulnerability to bacterial infection. Gas6-deficient mice displayed reduced mean blood pressure, exaggerated cytokine release, altered early leukocyte dynamics, and enhanced macrophage activation. Mechanistically, macrophages secreted Gas6 in response to LPS, and Gas6 acted as an autocrine negative feedback signal limiting inflammatory cytokine production, at least in part through regulation of TLR signaling.

Although Gas6 is strongly involved in immune regulation during sepsis, our data do not support a major direct role in coagulation activation. While circulating Gas6 and soluble Mer levels in septic patients were associated with sepsis-induced coagulopathy and impaired thrombin generation, this likely reflects disease severity rather than a causal effect on the coagulation. Consistently, in a murine endotoxemia model, Gas6 deficiency resulted in only minor changes in coagulation markers and did not lead to disseminated intravascular coagulation, suggesting that Gas6 primarily reflects systemic inflammation rather than directly driving coagulation abnormalities.

## Introduction

Sepsis is a life-threatening condition characterized by organ dysfunction resulting from a dysregulated host response to infection^1^, and it accounts for approximately 1 in 5 deaths worldwide^2^. In recognition of its substantial global burden, the World Health Organization has identified sepsis as a major public health priority^3^. Without early diagnosis and prompt management, sepsis can rapidly progress to septic shock, multiple organ failure, and death^4,5^.

In sepsis, simultaneous activation of pro- and anti-inflammatory pathways leads to the systemic release of cytokines, inflammatory mediators, and pathogen-associated molecules, with subsequent activation of the coagulation, fibrinolytic, and complement systems^6-10^. The process is initiated by the recognition of pathogen- and damage-associated molecular patterns (PAMPs and DAMPs), which bind to Toll-like receptors on monocytes and antigen-presenting cells. This interaction triggers transcriptional programs involved in inflammation, adaptive immunity, and cellular metabolism, driving the clinical syndrome. The resulting dysregulated inflammatory response contributes to progressive tissue injury and ultimately multi-organ dysfunction^11^. In later stages, a concurrent immunosuppressive state—characterized by reduced expression of activating surface molecules, increased immune-cell apoptosis, and T-cell exhaustion—predisposes patients to secondary infections, opportunistic pathogens, and viral reactivation^5,12-15^.

Over the past three decades, numerous clinical trials have been initiated based on promising results from animal models of sepsis; however, these findings have largely failed to translate into effective therapies in human patients^16^. As a result, no specific curative therapy for sepsis is currently available, underscoring the urgent need for new treatment strategies. Therefore, understanding sepsis at the molecular and cellular levels is mandatory along with the use of improved preclinical models closely associated with *ex vivo* and *in vitro* studies on human cells and, when also possible, tissues, in order to improve the quality of the translational research on sepsis^16,17^.

*Gas6* gene was originally identified as one of the genes upregulated in fibroblasts during growth arrest^18^. Gas6 protein belongs to the vitamin K-dependent protein family^19^. It has a significant homology with anticoagulant protein S, both in modular organization and amino-acid residue composition^19^. Like all vitamin K-dependent proteins, Gas6 binds to anionic phospholipids via its calcium-bound Gla domain^20^. Through its SHBG-like domain, Gas6 binds to receptors present on the cell surface^21^. Gas6 is expressed by a variety of cell types, including endothelial cells^19^, vascular smooth muscle cells^22^, and bone marrow cells^23^. Consistent with this wide distribution, Gas6 has been implicated in various crucial cell functions including reversible cell growth arrest^19^, survival^24^, proliferation^24-28^, cell adhesion^29-31^ and cell migration^31,32^.

Gas6 and protein S activate the TAM family of receptor tyrosine kinases (Tyro3, Axl, and Mer), triggering signaling pathways involved in diverse biological functions ^33^. Gas6 is a ligand for all three TAM receptors, with the highest affinity for Axl, whereas protein S binds only Tyro3 and Mer^34,35^.

Elevated plasma Gas6 levels have been consistently reported in severe sepsis and correlate with disease severity and organ dysfunction, suggesting potential prognostic value.^36-41^,^36,37,39^ However, reanalysis of the Need-Speed trial by Gavelli et al. found no association between admission levels of Gas6 or soluble Mer (sMer) and 7- or 30-day mortality.^42^ Instead, both markers predicted thrombocytopenia and sepsis-induced coagulopathy, with higher levels observed in affected patients. Experimental studies further support a protective role for Gas6, showing reduced neutrophil migration, attenuation of lung injury, and preservation of endothelial barrier integrity in sepsis models^43^ ^44^ ^45^. Consistent with these findings, Gas6 administration in mouse models of sepsis-related kidney injury, lung injury, and peritonitis improved vitality and limited organ damage, particularly when combined with antibiotic^43,44,46,47^.

Gas6 is secreted by endothelial cells and, in mice, by activated monocytes and platelets. Because monocytes and endothelial cells are key responders during inflammation, they are likely major sources of circulating Gas6 in sepsis. This is supported by its correlation with plasma thrombomodulin, a marker of endothelial injury, and by the association between Gas6 mRNA levels and monocyte counts in septic patients but not in noninfectious organ failure^36^, indicating infection-specific production. In addition, Gas6 mRNA levels in whole blood correlate with monocyte counts in septic patients, but not in patients with organ-failure without infection, indicating that Gas6 secretion occurs specifically during infection^36^. Beyond its release, Gas6 also amplifies inflammatory responses by promoting endothelial activation, facilitating the adhesion of platelets and leukocytes to the endothelium, and enhancing leukocyte extravasation and thrombosis^48^.

Mer deficient (*Mer^-/-^*) mice are hypersensitive to endotoxemia, showing increased mortality and organ damage driven by excessive TNF-α production ^49,50^. Similarly, lipopolysaccharide (LPS)-stimulated antigen-presenting cells from TAM-deficient mice display heightened activation, with increased expression of MHC class II, CD86, IL-12, TNF-α, and NF-κB. Endothelial ICAM-1 is also upregulated, promoting lymphocyte adhesion^50^. Axl expression is induced in monocytes by IFN-γ, and mice overexpressing Axl in myeloid cells produce elevated TNF-α, even in response to minimal LPS stimulation^51^.

Axl activation also triggers anti-inflammatory feedback by inducing SOCS1 and SOCS3 through the type I interferon receptor–STAT1 pathway. These molecules act as negative regulators of cytokine signaling and help integrate Axl into broader signaling complexes^52^ ^53^.

Based on previous studies that report increased Gas6 plasma levels in severe sepsis^36-39^, we decided to investigate Gas6, as a likely candidate for sepsis treatment.

The findings of this study support a role for Gas6 signaling through its receptors in shaping the host response to infection and endotoxemia by modulating innate immunity. The observed increase in plasma Gas6 in healthy volunteers during endotoxin challenge is consistent with experimental data in mice and suggests that endogenous Gas6 may act as a protective factor in severe sepsis, helping to limit macrophage-driven inflammation after the initial endotoxin-induced activation. Moreover, administration of exogenous Gas6 protects mice from endotoxemia, likely by binding soluble Axl and thereby increasing the availability of free Gas6 to engage cellular TAM receptors and regulate the systemic innate immune response.

## Material and methods

### Human Samples

Human samples were obtained from two independent cohorts: severe sepsis patients, patients with septic shock, non-infectious inflammatory and healthy controls recruited in Geneva in 2007 and septic shock patients and healthy volunteers enrolled in Bern between 2023 and 2025 (REDUCE study, control arm only (NCT04931485)).

#### Sepsis cohort and controls (Geneva, 2007)

Twenty-nine adult patients with severe sepsis or septic shock (diagnosed according to consensus cirteria in ref^54^ were prospectively enrolled on ICU admission at Geneva University Hospitals, in 2007. Heparinised plasma was collected within 24 h of admission and stored at −80 °C. Additional plasma samples were obtained from 13 healthy subjects and 18 patients with acute exacerbation of rheumatoid arthritis as healthy and non-infectious inflammatory controls. The study protocol was approved by the institutional ethics committee and written informed consent was obtained from all participants or their legal representatives.

#### Bern REDUCE cohort (2023–2025)

Patients were enrolled between 2023 and 2025 as part of the REDUCE trial at Inselspital, Bern (NCT04931485). REDUCE was a randomised controlled multicentre trial evaluating a fluid management protocol in patients with septic shock according to the Sepsis-3 criteria^1^. For the present biomarker analyses, only control-arm patients (i.e. receiving standard of care) were included, and additional blood samples were taken. The REDUCE trial and associated biomarker sub-study were approved by the Ethics Committee of the Canton of Bern (BASEC-Nr. 2020-02197), and all participants or their legal representatives provided written informed consent.

Third cohort:

Healthy volunteers (28–35 years) were recruited among staff at the University of Bern through 2022-2025. Inclusion criteria were self-reported good health, no regular prescription medication, and no intake of non-steroidal anti-inflammatory drugs (including ibuprofen) or aspirin during the preceding 2 weeks.

### Mice

*Gas6^+/+^* and *Gas6^-/-^* (***B6.129S1-Gas6^tm1Pec^***) mice used in most experiments were on a C57BL/6J background. Mice were bred in the central animal facility of the University of Bern and used at 8 – 18 weeks of age. They were housed in ventilated cages under a 12 h light-dark cycle at 20 ± 2 °C with a relative humidity of 50 ± 10% and had ad libitum access to standard food and water. Age-, sex- and strain-matched *Gas6^+/+^* controls were used in all experiments. All animal procedures complied with Swiss regulations for the care and use of laboratory animals and the ARRIVE guidelines. Experiments were approved by the Animal Care Committee of the Canton of Bern.

Other experiments (as indicated in the corresponding figure legends) used *Gas6^+/+^*, *Gas6^+/-^* and *Gas6^-/-^* mice derived from the original colony with a mixed genetic background of 50% 129/Sv and 50% Swiss, as previously described^55^. BALB/cAnNCrl *Gas6^-/-^* mice were backcrossed for more than ten generations on a BALB/cAnNCrl background. *Tyro3^-/-^*, *Axl^-/-^* and *Mer^-/-^* mice were progeny of the original colony on a 50% 129/Sv x 50% C57BL/6 background, and *Gas6^+/+^* control mice shared the same genetic background^56^.

### Murine bacterial sepsis model

In the first model, *Escherichia coli* O18 strain (kindly provided by A.S. Cross)^57^ was cultured in brain heart infusion medium (Becton Dickinson, Franklin Lakes, NJ, USA) for 3 – 4 h at 37 °C until an optical density at 620 nm of 0.4 was reached, corresponding to approximately 3 x 10^8^ CFU/mL. BALB/cAnNCrl *Gas6^+/+^* and *Gas6^-/-^* mice were injected intraperitoneally with 1.5 x 10^4^ to 2.5 x 10^5^ CFU in the appropriate volume. The inoculum was plated immediately after injection on blood agar plates to determine the viable bacterial counts. In the CLP model, mice were anesthetized with halothane, and a midline laparotomy was performed to expose the cecum. The cecum was ligated as close to the ileocecal junction as possible and perforated twice with a 19-gauge needle, then gently squeezed to extrude a small amount of faecal material before being returned to the peritoneal cavity. The abdominal incision was closed with wound clips. Sham-operated mice underwent laparotomy and exteriorization of the cecum, which was then returned to the peritoneal cavity without ligation or puncture. After surgery, and every 12 h thereafter, mice received buprenorphine (Temgesic, Essex Chemie, Lucerne, Switzerland; 0.05 mg/kg, subcutaneously) for analgesia.

### Murine sepsis score (MSS)

Clinical severity was assessed using a predefined murine sepsis score sheet, which combined general appearance, behaviour and physiological parameters into a cumulative score. The following variables were scored from 0 (normal) to 3 (severely affected): fur aspect (actively grooming to piloerection), activity (normal to no activity when disturbed or stimulated), posture (normal to hunched with no movement), respiration (normal to severe dyspnoea), facial grimace (open ears to orbital tightening with cheek bulge and flat ears), body weight loss (0 – 5 % to >15 %) and decrease in body temperature (<0 °C to <-3 °C relative to baseline). Mice were monitored and scored over time. The cumulative score guided refinement measures and humane endpoints. A cumulative score of 1 – 4 triggered increased monitoring (at least twice daily) and provision of critical care rodent diet when body weight loss reached 5 – 10 %, whereas a score of 5 – 8 prompted administration of buprenorphine 0.1 mg/kg body weight subcutaneously (Bupaq® 0.3 mg/mL, Streuli, Tiergesundheit AG) and intensified monitoring up to three times daily. A cumulative score greater than 8, a score of 3 for dyspnoea, or persistent high scores in combination with ≥20 % body weight loss mandated euthanasia, in accordance with the approved animal experiment protocol.

### Low dose endotoxemia in healthy volunteers

Eight healthy young men^58^ were enrolled after approval by the institutional ethics committee and written consent. Inclusion criteria were good health and no medication. A complete history, physical examination and 12-lead electrocardiography were performed. Peripheral venous blood was collected into vacutainer tubes containing 3.2% (0.109 M) sodium citrate anticoagulant at a 1:9 (v/v) ratio prior to 2 ng/kg Escherichia coli LPS administration (baseline) and at predefined timepoints following intravenous injection to capture dynamic inflammatory and hemostatic responses during the controlled human endotoxemia model (standard dose used to induce a controlled human endotoxemia model in healthy volunteers, as detailed in the original study^58^. Participants provided written informed consent.

### Murine endotoxemia model

Mice were injected intraperitoneally with LPS from *Escherichia coli* O55:B5 (Sigma-Aldrich) at doses of 10, 25, 40 or 50 mg/kg and monitored for up to 72 h for LPS-induced lethality.

### Plasma preparation

For the mouse thrombin-antithrombin (TAT) assay, blood was collected into tubes containing 3.2% (w/v) sodium citrate. Platelet-poor plasma (PPP) was prepared by centrifugation of citrated whole blood at 2,500 x *g* for 10 min at room temperature (22 °C).

For thrombin generation (TG) assays, citrated blood was centrifuged twice at 2,500 x *g* for 10 min at room temperature.

For biomarker quantification (TNF-α, IL-6, Gas6, sAxl, sMer, and sTyro3), either citrated or lithium heparinized blood was centrifuged once at 10,000 x *g* for 10 min at room temperature.

Human plasma samples were prepared similarly by centrifuging citrated blood twice at 2,500 x *g* for 15 min at room temperature. All plasma samples were snap-frozen on dry ice, stored at −80 °C.

### ELISA

TAT complexes were measured in duplicate in mouse plasma using a commercial ELISA kit (Enzygnost TAT micro, Siemens Healthineers; cat. no. 23-044-704) according to the manufacturer’s instructions. For murine biomarkers, Gas6, sAxl, sMer, sTyro3, TNF-α, IL-6 and IFN-γ were quantified by sandwich ELISA following the manufacturer’s instructions (Mouse DuoSet ELISA kits, R&D Systems: Gas6, DY986; Axl, DY854; Mer, DY591; Tyro3/Dtk, DY759; IL-6, DY406; TNF-α, DY410; IFN-γ, DY485). For human samples, Gas6, sAxl, sMer and sTyro3 were measured using Human DuoSet ELISA kits (R&D Systems: Axl, DY154; Gas6, DY885B; Mer, DY6488; Tyro3/Dtk, DY859) according to the manufacturer’s protocols, and nitrate plus nitrite concentrations were determined with a colorimetric assay (Cayman Chemical) following the manufacturer’s instructions.

### Recombinant proteins

For in vitro assays, recombinant Gas6 (rGAS6) was expressed and purified from HEK293 cells transfected with an epitope-tagged Gas6 cDNA, as previously described^59^.

For *in vivo* injection experiments, recombinant full-length mouse Gas6 protein was purchased from R&D Systems (Recombinant Mouse Gas6, CF, cat. no. 8310-GS), was diluted in PBS, and injected 10 ug/mouse via tail vein.

### Gas6-sAxl complex with ELISA

For complex evaluation in human samples, Gas6-sAxl complexes were quantified using Human DuoSet ELISA kits (R&D Systems: Axl, DY154; Gas6, DY885B) according to the manufacturer’s protocols with modifications. We used Gas6 Capture Ab (400 ng/mL final concentration in well) and Axl Detection Ab (0.63 μg/mL final concentration in well).

For complex evaluation in mouse samples, Gas6-sAxl, were quantified sandwich ELISA following the manufacturer’s instructions (Mouse DuoSet ELISA kits, R&D Systems: Gas6, DY986; Axl, DY854) and using Gas6 Capture Ab and Axl Detection Ab. As washing buffer 1x TBS pH 7.4, 1.3mM CaCl2 and 0.1% Tween 20, was used.

### Arterial blood pressure measurements

Male *Gas6^+/+^* and *Gas6^-/-^* mice were i.p. injected with 40 mg/kg *E. coli* LPS or an equivalent volume of isotonic saline. After 6 hours, arterial blood pressure was measured inserting a 1.4 F micro-tip pressure catheter (Millar Instruments) into the right carotid artery of anesthetized mice (ketamine 80 mg/kg and xylazine 10 mg/kg). Signals were recorded with a Powerlab/4SP A/D converter (ADInstruments).

### Mouse protein cytokine array

Mouse plasma samples or macrophage supernatants were analysed using a mouse cytokine protein array detecting 62 cytokines (RayBiotech) (**Table 1**). Membranes were processed according to the manufacturer’s instructions: after blocking in the supplied blocking buffer, 1 mL of pooled plasma (from 8 mice) or macrophage supernatant was added per membrane and incubated at room temperature for 2 h. Arrays were then washed, incubated with detection reagents, and chemiluminescent signals were recorded and quantified using a VerSaDoc Imaging System (Bio-Rad), and spot intensities were graphed for downstream analysis.

**Table 1.**
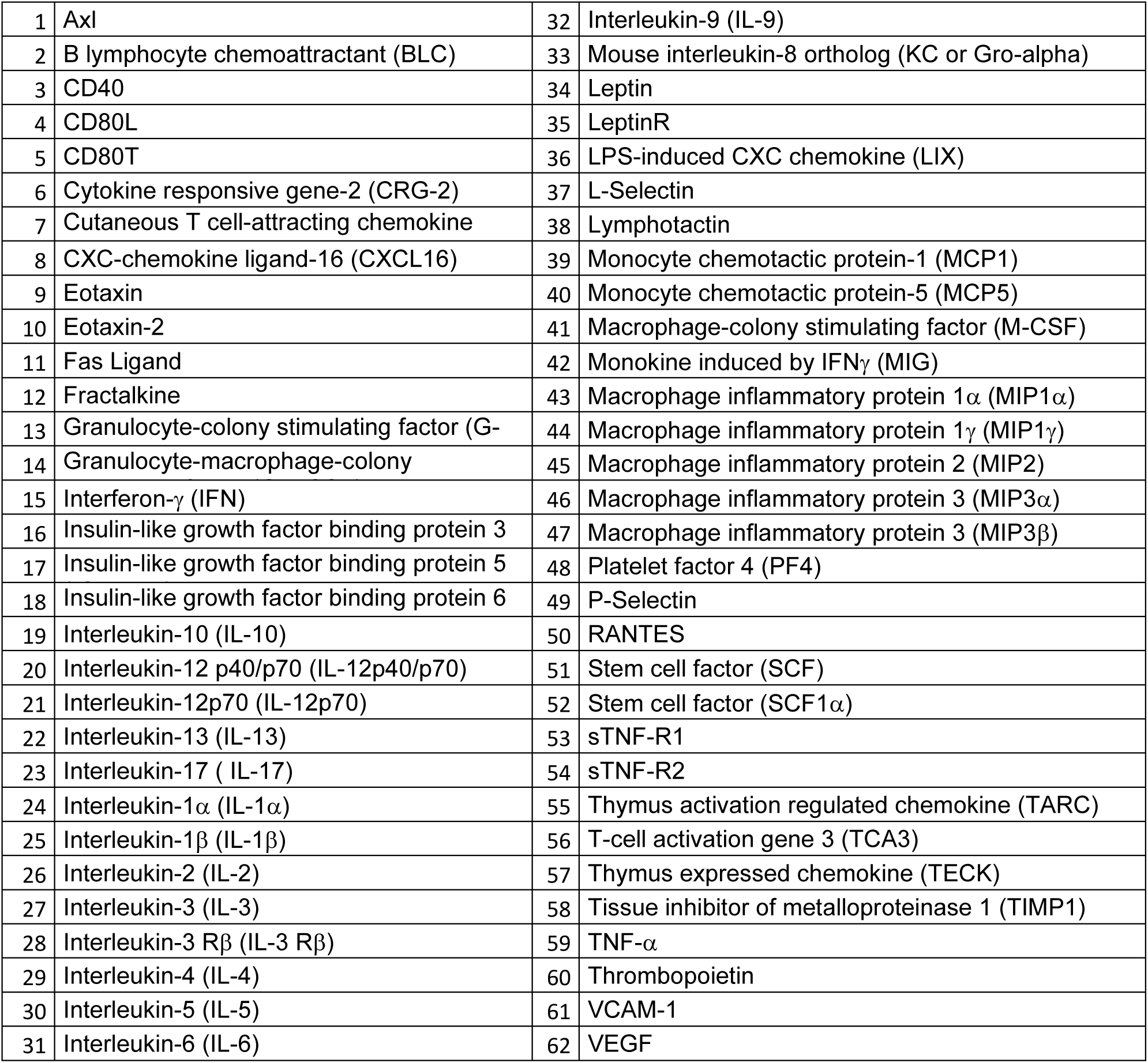
Cytokine array list.

### Complete Blood Counts

Complete blood counts were measured using an automated veterinary haematology analyser (ProCyte Dx Hematology Analyzer, IDEXX Laboratories). Reagent and stain kits (IDEXX ProCyte Dx Reagent Kit, 388401; IDEXX ProCyte Dx Stain Pack, 388400) were routinely replaced according to the manufacturer’s instructions, and 30 µL of whole blood was used per measurement.

### Bone marrow-derived macrophages (BMDMs) culture

BMDMs were differentiated from mouse bone marrow cells in complete macrophage medium containing 30% (v/v) L929-conditioned supernatant and 70% base medium (DMEM with GlutaMAX, 10% HI FCS, 100 IU/mL penicillin, 100 μg/mL streptomycin). Bone marrow cells from mouse femurs, were seeded in non-treated tissue culture dishes and cultured for 7 days at 37 °C, 5% CO2, with medium changes every 3-4 days using fresh L929-conditioned medium. At day 7, adherent cells with macrophage morphology were detached using Accutase (Biolegend, cat. no. 423201) and used for downstream experiments.

### Mass cytometry peripheral blood mononuclear cells (PBMCs)

A novel protocol for immunophenotyping of PBMCs in mouse blood was established.

Blood was collected from mice vena cava into syringes preloaded with sodium heparin (final concentration 40 IU/mL in 500 µL blood) and kept at room temperature. Red blood cells were lysed by adding 5 mL ACK lysis buffer (Gibco, A10492-01) in a 50 mL polypropylene tube, gently inverting three times and incubating for 5 min at room temperature. Lysis was stopped by adding 10 mL ice-cold PBS. Cells were pelleted (350 x *g*, 10 min, 4 °C), the supernatant was removed, and – if residual erythrocytes were visible – the lysis step was repeated with 2.5 mL ACK buffer followed by dilution in 5 mL ice-cold PBS.

After lysis, cells were washed with 10 mL PBS (350 x *g*, 10 min, 4 °C) and the pellet was resuspended in 250 µL ice-cold cell staining buffer (CSB; DPBS supplemented with 2 mL 0.5 M EDTA and 3 mL FBS per 500 mL). Cells were counted, washed once more in 5 mL CSB (350 x *g*, 5 min, 4 °C) and resuspended in 100 µL CSB. For barcoding and viability staining, 1 µL of each anti-mouse CD45 metal barcode antibody was added to wells of a 96-well U-bottom plate, and 100 µL of the cell suspension was added per well; samples were incubated for 1 h at 4 °C on ice with gentle mixing for 30 min. Cells were then washed three times with CSB (350 x *g*, 3 min, 4 °C), pooled, and incubated with cisplatin (diluted 1:10,000 in protein-free PBS; 200 µL per 2 × 10^6^ cells) for 5 min at room temperature, followed by two CSB washes.

For Fc receptor blocking and surface staining, cells were incubated with mouse Fc block (TruStain FcX; 100 µL per 2 × 10^6^ cells, 1:200 in CSB) for 10 min at room temperature without subsequent washing. A surface antibody cocktail (100 µL per 2 × 10^6^ cells in CSB) was then added and cells were incubated for 30 min at room temperature with gentle mixing after 15 min. Cells were washed twice with CSB (240 µL, 350 x *g*, 3 min, 4 °C).

For final fixation, pellets were gently resuspended in 200 µL per 2 × 10^6^ cells of freshly prepared 4 % formaldehyde (from 16 % stock diluted in Maxpar PBS) and incubated for 10 min at room temperature before centrifugation (800 x *g*, 3 min, 4 °C) without further washing.

For DNA intercalation and acquisition, cells were incubated with iridium intercalator diluted 1:5,000 in Maxpar Fix and Perm Buffer (100 – 200 µL per 2 x 10⁶ cells) for 1 h at room temperature. When samples were not acquired immediately, plates were sealed and stored at 4 °C overnight or frozen at −80 °C. On the day of acquisition, cells were thawed on ice, filtered and acquired on a Helios™ CyTOF instrument using standard settings for murine PBMC panels. The surface antibody panel was designed and titrated in collaboration with Deborah Stroka laboratory (Department of Visceral Surgery and Medicine, Inselspital, Bern University Hospital and DBMR, University of Bern) with metal tags, targets, clones, and final dilutions as detailed in **Table 2**.

**Table 2.** List of Antibodies used for PBMCs staining by mass cytometry protocol.

| **Surface Antibody Cocktail** |  |  |  |
|---|---|---|---|
| **Metal Tag** | **Target** | **Antibody Clone** | **Titration** |
| Dy163 | CD86 | PO3 | 100 |
| Yb171 | CD80 | 16-10A1 | 100 |
| Eu151 | CD64 | X54-5/7.1 | 100 |
| Er167 | CX3CR1 | SA011F11 | 200 |
| Yb176 | CD192 | 475301 | 200 |
| Sm154 | CD40 | HM40-3 | 200 |
| Dy164 | CD49b | HMa2 | 100 |
| Sm149 | CD19 | 6D5 | 400 |
| Sm152 | CD3e | 145-2C11 | 200 |
| Nd142 | CD11c | N418 | 800 |
| Nd150 | Ly6C | HK1.4 | 800 |
| Yb174 | MHC II I-A/I-E | M5/114.15.2 | 800 |
| Nd144 | CD115 | AFS98 | 800 |
| Nd148 | CD11b | M1/70 | 1600 |
| Er170 | CD161 | PK136 | 400 |
| Pr141 | Ly6G | 1A8 | 1600 |
| Er168 | CD8a | 53-6.7 | 1600 |
| Yb172 | CD4 | RM4-5 | 1600 |

### Intravital microscopy of mesenteric venules

Mice (8–9 week-old males and females) received an intraperitoneal injection of LPS (50 mg/kg) and were imaged 3 h later under MMF anaesthesia (0.05 mg/kg fentanyl, 5 mg/kg midazolam (Dormicum) and 0.5 mg/kg medetomidine; 3 µL/g body weight i.p. in 0.9% NaCl), prepared fresh on the day of the experiment.

A tail-vein catheter (polyethylene tubing, inner/outer diameter 0.11 × 0.24 mm) was inserted. Anaesthetised mice were placed on a heated surgical pad, eyes were protected with ophthalmic gel, and the tail-vein catheter was inserted and flushed with 0.9% NaCl; anaesthesia was supplemented via the catheter as needed until pedal reflexes were absent. A midline laparotomy (∼2 cm) was performed, the small intestine was gently exteriorised, and the mesentery was spread without tension on a temperature-controlled intravital microscopy stage; the gut was stabilised with NaCl-soaked Kimtech paper while leaving the mesenteric vessels uncovered.

Fluorescently labelled antibodies against P-selectin (clone Psel.KO2.3; PE; eBioscience™, Invitrogen™, cat. no. 15238579), CD31 (clone 390; Alexa Fluor 647; BioLegend, cat. no. 102416) and CD45 (clone 30-F11; FITC; eBioscience™, Invitrogen™, cat. no. 15238089) were injected through the tail-vein catheter (10 µL of each antibody), followed by a 50 µL flush of 0.9% NaCl. Intravital imaging was performed on a Leica IVM system controlled by LAS X software using a 25x objective and multiphoton plus confocal laser lines (405, 488, 561 and 638 nm), with detectors and laser shutters adjusted accordingly. Mesenteric post-capillary venules with diameters of approximately 150–200 µm were selected, and images were acquired in Xyt mode at 512 × 512 pixels using the resonant scanner (8,000 Hz, zoom 1.25, line average 6, rotation 0); time series of up to 30 min were recorded for quantification of leukocyte rolling and adhesion. In septic mice, images were acquired every 1 s, whereas in control mice images were acquired every 2 s to account for differences in blood-flow velocity. For euthanasia, mice received an intraperitoneal overdose of pentobarbital (150 mg/kg body weight; Eskonarkon® 300 mg/ml diluted 1:20 in 0.9% NaCl).

### Whole blood thrombin generation assay

Whole blood thrombin generation was performed as described by Wan et al.^60^, with the following modifications. Citrated whole blood (3.2% sodium citrate, 9:1 blood:anticoagulant) was collected through vena cava puncture for mice to minimize TF contamination, venipuncture for humans and assayed within 1 h of collection. The fluorogenic substrate Z-Gly-Gly-Arg-aminomethylcoumarin (ZGGR-AMC) was used, and measurements were carried out in U-bottom 96-well microplates (Immulon 2HB, cat. no. 735-0464). Human tissue factor was provided as Innovin (Dade Behring, Marburg, Germany), and fluorescence was recorded for 60–80 min at 37 °C on a Fluoroskan Ascent microplate fluorometer (Thermo Scientific) with excitation/emission wavelengths of 355/460 nm controlled by the Fluoroskan Ascent software. A mixture 120 uL blood, 40 uL substrate (final concentration 0.41 mM), 80 uL Calibrator (final concentration 100 nM) or TF (final concentration 0.5-1 pM)/CaCl2 (final concentration 9 mM) was prepared. A part of this mixture was separated in equal volumes of triplicate detection wells (65 uL per well). The assay was performed similarly for mouse and human citrated whole blood, with no changes in reagents, buffers or plate setup. Substrate was prediluted in BSA buffer containing 20 mM Hepes, 6% BSA, pH 7.35, and TF/CACl2 was prediluted in BSA buffer 20 mM Hepes, 140 mM NaCl, 0.5% BSA, pH 7.35. Key TG outputs were recorded: lag time (min), peak height (nM), time-to-peak (min), endogenous thrombin potential (ETP, nM·min), start tail, TG (thrombin (nM) mean of triplicates per each individual sample.

### Thrombin generation assay

Thrombin generation was measured in murine using a calibrated automated thrombography (CAT) approach in U-bottom 96-well microplates (Immulon 2HB, Thermo Fisher Scientific, Waltham, MA, USA) at 37 °C. Mouse PPP samples were thawed in a 37 °C water bath for 10 min and diluted 1:3 in Buffer A (20 mM HEPES, 140 mM NaCl, 5 mg/mL BSA, pH 7.4) before use, whereas human PPP samples were used undiluted. Low tissue factor trigger (∼1 pM TF, PPP-reagent Low, Stago, cat. no. 86194), a microparticle-rich trigger (MP-reagent, Stago, cat. no. 86222), a thrombin calibrator (Stago, cat. no. 86192) and thrombomodulin (Synapse) were reconstituted with distilled water according to the manufacturer’s instructions.

For each condition, samples were run at least in duplicate. In each well, 10 µL thrombin calibrator or 10 µL PPP-reagent or 10 µL MP-reagent or with 2 nM TM were added as appropriate, followed by 40 µL PPP (prediluted mouse PPP) and 40 µL fluorogenic substrate solution (diluted in 1.6 mL Fluo-Buffer; FluCa kit, Stago; cat. no. 86197) to start the reaction. Fluorescence was recorded on a Fluoroskan Ascent fluorescence plate reader (Thermo Scientific, Waltham, MA, USA) using 390 nm excitation and 460 nm emission, with readings every 20 s for 90 min, under Thrombinoscope software control (version 5.0.0.742).

### Halo fluorescence fibrinolysis (HoFF)

HoFF test was performed as previously described (Liu et al., 2025). Briefly, citrated platelet-poor plasma from mice was mixed with Alexa Fluor 488–conjugated fibrinogen and clotted with thrombin/CaCl₂ in 96-well flat-bottom plates to generate halo-shaped plasma clots, which were then overlaid with tissue plasminogen activator–containing buffer and monitored by microplate reader (Tecan) at 37 °C.

### Laboratory measurements

Routine clinical chemistry parameters including plasma fibrinogen, prothrombin time, creatinine, alanine aminotransferase, aspartate aminotransferase, were measured on an automated Sysmex CA-7000 coagulation analyser (Sysmex Digitana, Zentrum für Labormedizin, Inselspital, Bern, Switzerland) using standard methods.

### Western blotting for signal transduction and quantification of phosphorylated IκBα

For Western blotting, MBDMs were washed with cold PBS and lysed directly on plates at 4°C with Cell Lysis Buffer (Cell Signaling Technology) supplemented with Complete Mini protease inhibitors cocktail (Roche Diagnostics) and 1 mM NaF. Lysates were centrifuged 10 minutes at 17’000 x g, 4°C and then supernatants were kept at -80°C until use. Proteins were quantified using BCA Protein Assay Kit (Pierce Biotechnology) separated by SDS-PAGE and transferred onto nitrocellulose (Amersham). Phosphorylation of Akt (at Ser473), ERK1/2 (Thr202/Tyr204), p38 (Thr180, Tyr182) and JNK (Thr183/185) was determined using 1:1000 dilutions of rabbit antibodies (Cell Signaling Technologies) incubated overnight at 4°C. For total protein control, membranes were stripped in 62.5 mM Tris-HCl pH=6.8, 2% SDS, 100 μM β-mercaptoethanol for 30 minutes at 50°C, and reprobed with antibodies (Cell Signaling Technologies) directed against the total form of the protein (dilution 1:1000). All primary antibodies were detected by HRP-conjugated swine anti-rabbit antibodies (dilution 1:3000, DakoCytomation). Membranes were incubated with ECL (Amersham) and exposed to Hyperfilm (Amersham). Phosphorylated and total IκBα were quantified in macrophages by an ELISA kit (PathScan ELISA kits, Cell Signaling Technology) according to manufacturer’s instructions.

### SOCS1 and SOCS3 quantification by qPCR

Gene expression of SOCS1 and SOCS3 was quantified by real-time qPCR, relative to the expression of GAPDH, using an ABI PRISM 7000 (Applied Biosystems) real-time cycler and the following primers: GAPDH forward, 5′-CAACGGGAAGCCCATCAC-3′; reverse, 5′-CGGCCTCACCCCATTTG-3′; SOCS1 forward, 5′-CCGTGGGTCGCGAGAAC-3′; reverse, 5′-AACTCAGGTAGTCACGGAGTACCG-3′; SOCS3 forward, 5′-TCCCATGCCGCTCACAG-3′; reverse, 5′-ACAGGACCAGTTCCAGGTAATTG-3′.

Each reaction contained 4 μl SYBR Green PCR Master Mix (Applied Biosystems), 50 nM primer pair, 5 μl diluted cDNA (obtained with SuperScript II Reverse Transcriptase (Invitrogen) according to manufacturer instructions) and water to a final volume of 20 μl. PCR parameters were as follows: initial denaturation at 95°C for 10 min followed by 40 cycles of 15 s at 60°C. The relative gene expression levels in each sample were determined using the comparative deltadeltaCt method, and the GAPDH gene as the endogenous control.

### Statistical analyses

Statistical analyses were performed using GraphPad Prism version 10.2.3 (GraphPad Software, San Diego, CA, USA). Data are presented as mean ± standard error of the mean (SEM). Normality of distributions was assessed using the Shapiro–Wilk test (α = 0.05); parametric tests were applied to data meeting normality assumptions, with non-parametric alternatives used otherwise. For multiple-group comparisons, one-way ANOVA was used for single-factor experiments, followed by post hoc Sidak’s or Holm–Sidak’s tests for pairwise interactions. Two-way ANOVA was employed for experiments with two independent variables (e.g., genotype and treatment), with post hoc Sidak’s or Holm–Sidak’s tests for pairwise interactions. For two-group comparisons, unpaired two-tailed t-tests were performed assuming equal variances. Non-parametric equivalents included Kruskal–Wallis one-way ANOVA with Dunn’s post hoc test for >2 groups and Mann–Whitney U tests for two groups.

Statistical significance was accepted at P < 0.05. (P > 0.05, not significant, ns; *P < 0.05; **P < 0.01; ***P < 0.001; ****P < 0.0001).

## Results

### Plasma Gas6 levels increase during infection and endotoxemia in humans and mice and are associated with adverse outcomes in sepsis

Elevated plasma Gas6 levels have been reported in patients with sepsis and are associated with poorer outcomes^36-39^. To confirm and extend these findings, we compared circulating Gas6 concentrations at the time of severe sepsis diagnosis (n = 29) with those measured in healthy volunteers (n = 13) and in patients with rheumatoid arthritis (n = 18) as an inflammatory disease control. The characteristics of the patients included in Study 1are provided in **Table 3**. Gas6 levels tended to be higher in patients with rheumatoid arthritis than in healthy volunteers, although this difference did not reach statistical significance. In contrast, Gas6 concentrations were significantly elevated in septic patients relative to healthy volunteers, indicating that Gas6 increases in the setting of systemic inflammation (**Fig. 1A**). Notably, Gas6 levels were also significantly higher in septic patients than in those with rheumatoid arthritis, suggesting a stronger induction in the context of severe infection. In a second cohort comprising patients with septic shock (Study 2, Bern cohort, **Table 4**), Gas6 concentrations were already elevated within 24 hours of hospital admission and remained high for at least 7 days (**Fig. 1B**). This observation is consistent with our previous report showing sustained elevation of Gas6 for up to 40 days after hospital admission, with a gradual daily decrease of approximately 1.4%^39^. Furthermore, in Study 2, Gas6 concentrations positively correlated with SOFA scores (**Fig. 1C**), further supporting an association between Gas6 levels and disease severity.

**FIGURE 1:**
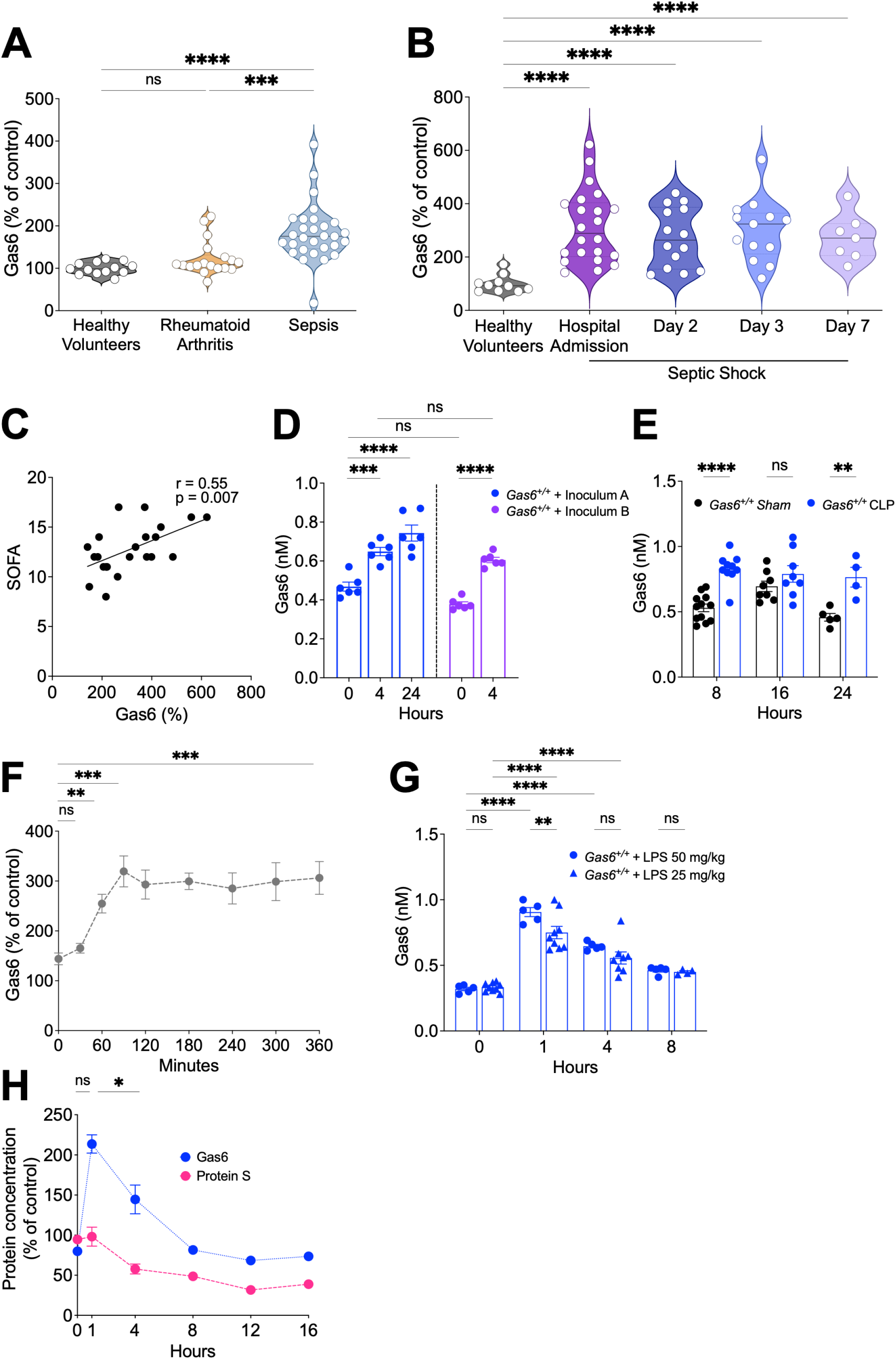
Plasma Gas6 levels increase during infection and endotoxemia in humans and mice and are associated with adverse outcomes in sepsis. (**A**), Gas6 plasma levels in healthy controls, patients with rheumatoid arthritis and septic patients at hospital admission. Kruskal-Wallis, correction: Dunnett’s multiple comparisons test (**B**), Gas6 plasma levels in healthy controls, and in patients with septic shock at hospital admission and at day 2, day 3, day 7. 2-way ANOVA, Dunnett’s multiple comparisons test (**C**), Spearman correlation and linear regression analyses assessing the relationship between plasma Gas6 levels and SOFA score in patients with septic shock. (**D**), *Gas6^+/+^* mice were inoculated intraperitoneally with the *E. coli* strain O18^57^ (inoculum A: 1.5*10^4^ CFU, inoculum B: 1.5*10^4^ to 2.5*10^5^ CFU) and plasma Gas6 levels in *Gas6^+/+^* mice were measured at different timepoints (n = 6). 2-way ANOVA, Dunnett’s multiple comparisons test (**E**), Gas6 plasma levels after 8, 16 and 24 h from CLP compared to sham-operated mice. 2-way ANOVA, Bonferroni’s multiple comparisons test (**F**), Gas6 levels in plasma of healthy volunteers (n=8) at steady-state (t0) and after 2 ng/kg LPS i.v. injection, repeated measures ANOVA and Bonferroni’s multiple comparison test. (**G**), Gas6 levels in plasma of *Gas6^+/+^* mice at steady-state and at different time points after LPS injection. Data from two independent experiments. (**H**), Gas6 and protein S levels in plasma of *Gas6^+/+^* mice at different time points after LPS (25 mg/kg) injection. ns, non-significant, * P < 0.05, **P < 0.01, ***P < 0.001,****P < 0.0001

**Table 3.**
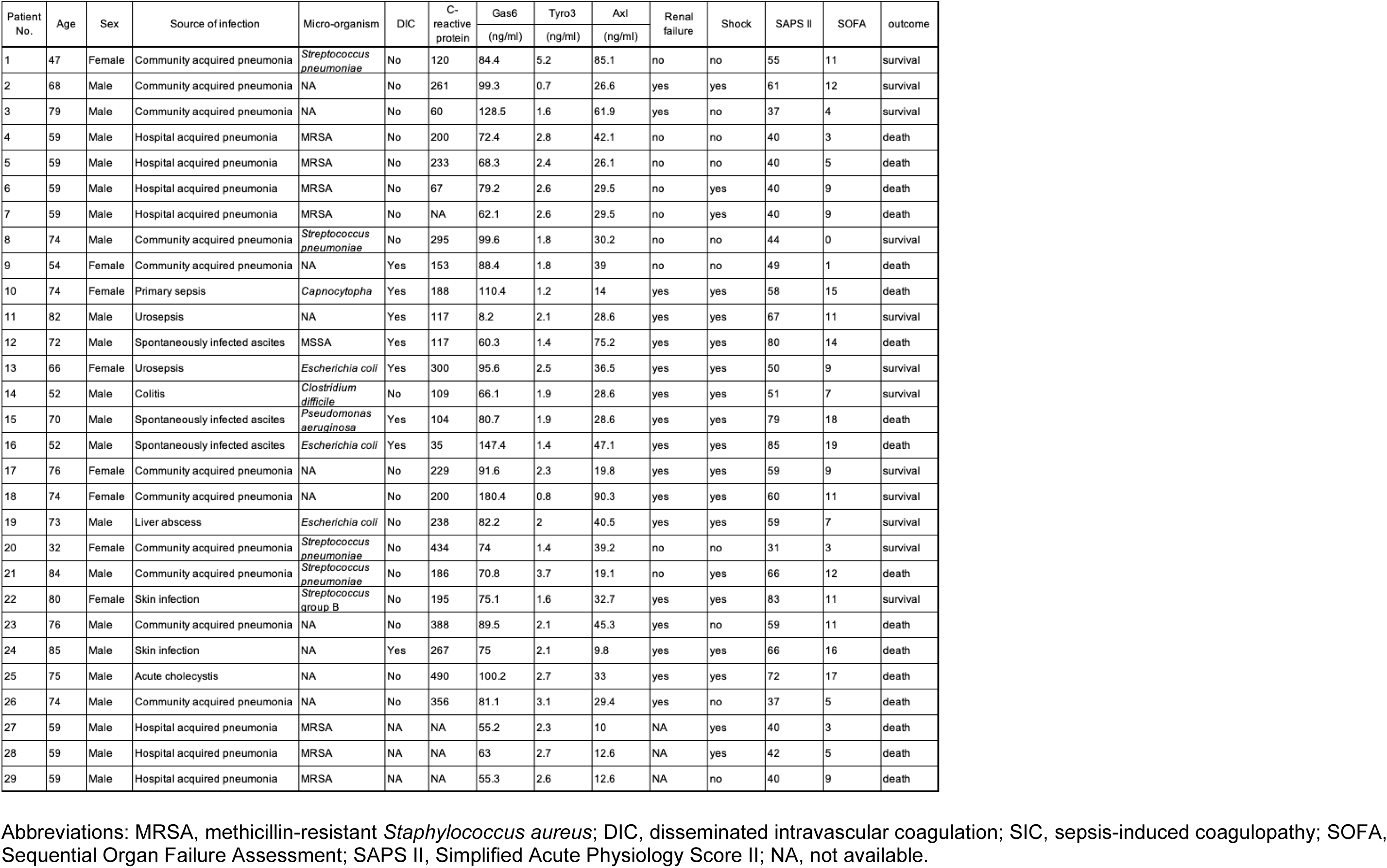
Patient characteristics (Study 1)

| Patient No. | Age | Sex | Source of infection | Micro-organism | DIC | C-reactive protein | Gas6 | Tyro3 | Axl | Renal failure | Shock | SAPS II | SOFA | outcome |
|---|---|---|---|---|---|---|---|---|---|---|---|---|---|---|
|  |  |  |  |  |  |  | (ng/ml) | (ng/ml) | (ng/ml) |  |  |  |  |  |
| 1 | 47 | Female | Community acquired pneumonia | *Streptococcus pneumoniae* | No | 120 | 84.4 | 5.2 | 85.1 | no | no | 55 | 11 | survival |
| 2 | 68 | Male | Community acquired pneumonia | NA | No | 261 | 99.3 | 0.7 | 26.6 | yes | yes | 61 | 12 | survival |
| 3 | 79 | Male | Community acquired pneumonia | NA | No | 60 | 128.5 | 1.6 | 61.9 | yes | no | 37 | 4 | survival |
| 4 | 59 | Male | Hospital acquired pneumonia | MRSA | No | 200 | 72.4 | 2.8 | 42.1 | no | no | 40 | 3 | death |
| 5 | 59 | Male | Hospital acquired pneumonia | MRSA | No | 233 | 68.3 | 2.4 | 26.1 | no | no | 40 | 5 | death |
| 6 | 59 | Male | Hospital acquired pneumonia | MRSA | No | 67 | 79.2 | 2.6 | 29.5 | no | yes | 40 | 9 | death |
| 7 | 59 | Male | Hospital acquired pneumonia | MRSA | No | NA | 62.1 | 2.6 | 29.5 | no | yes | 40 | 9 | death |
| 8 | 74 | Male | Community acquired pneumonia | *Streptococcus pneumoniae* | No | 295 | 99.6 | 1.8 | 30.2 | no | no | 44 | 0 | survival |
| 9 | 54 | Female | Community acquired pneumonia | NA | Yes | 153 | 88.4 | 1.8 | 39 | no | no | 49 | 1 | death |
| 10 | 74 | Female | Primary sepsis | *Capnocytopha* | Yes | 188 | 110.4 | 1.2 | 14 | yes | yes | 58 | 15 | death |
| 11 | 82 | Male | Urosepsis | NA | Yes | 117 | 8.2 | 2.1 | 28.6 | yes | yes | 67 | 11 | survival |
| 12 | 72 | Male | Spontaneously infected ascites | MSSA | Yes | 117 | 60.3 | 1.4 | 75.2 | yes | yes | 80 | 14 | death |
| 13 | 66 | Female | Urosepsis | *Escherichia coli* | Yes | 300 | 95.6 | 2.5 | 36.5 | yes | yes | 50 | 9 | survival |
| 14 | 52 | Male | Colitis | *Clostridium difficile* | No | 109 | 66.1 | 1.9 | 28.6 | yes | yes | 51 | 7 | survival |
| 15 | 70 | Male | Spontaneously infected ascites | *Pseudomonas aeruginosa* | Yes | 104 | 80.7 | 1.9 | 28.6 | yes | yes | 79 | 18 | death |
| 16 | 52 | Male | Spontaneously infected ascites | *Escherichia coli* | Yes | 35 | 147.4 | 1.4 | 47.1 | yes | yes | 85 | 19 | death |
| 17 | 76 | Female | Community acquired pneumonia | NA | No | 229 | 91.6 | 2.3 | 19.8 | yes | yes | 59 | 9 | survival |
| 18 | 74 | Female | Community acquired pneumonia | NA | No | 200 | 180.4 | 0.8 | 90.3 | yes | yes | 60 | 11 | survival |
| 19 | 73 | Male | Liver abscess | *Escherichia coli* | No | 238 | 82.2 | 2 | 40.5 | yes | yes | 59 | 7 | survival |
| 20 | 32 | Female | Community acquired pneumonia | *Streptococcus pneumoniae* | No | 434 | 74 | 1.4 | 39.2 | no | no | 31 | 3 | survival |
| 21 | 84 | Male | Community acquired pneumonia | *Streptococcus pneumoniae* | No | 186 | 70.8 | 3.7 | 19.1 | no | yes | 66 | 12 | death |
| 22 | 80 | Female | Skin infection | *Streptococcus group B* | No | 195 | 75.1 | 1.6 | 32.7 | yes | yes | 83 | 11 | survival |
| 23 | 76 | Male | Community acquired pneumonia | NA | No | 388 | 89.5 | 2.1 | 45.3 | yes | no | 59 | 11 | death |
| 24 | 85 | Male | Skin infection | NA | Yes | 267 | 75 | 2.1 | 9.8 | yes | yes | 66 | 16 | death |
| 25 | 75 | Male | Acute cholecystis | NA | No | 490 | 100.2 | 2.7 | 33 | yes | yes | 72 | 17 | death |
| 26 | 74 | Male | Community acquired pneumonia | NA | No | 356 | 81.1 | 3.1 | 29.4 | yes | no | 37 | 5 | death |
| 27 | 59 | Male | Hospital acquired pneumonia | MRSA | NA | NA | 55.2 | 2.3 | 10 | NA | yes | 40 | 3 | death |
| 28 | 59 | Male | Hospital acquired pneumonia | MRSA | NA | NA | 63 | 2.7 | 12.6 | NA | yes | 42 | 5 | death |
| 29 | 59 | Male | Hospital acquired pneumonia | MRSA | NA | NA | 55.3 | 2.6 | 12.6 | NA | no | 40 | 9 | death |

Abbreviations: MRSA, methicillin-resistant *Staphylococcus aureus*; DIC, disseminated intravascular coagulation; SIC, sepsis-induced coagulopathy; SOFA, Sequential Organ Failure Assessment; SAPS II, Simplified Acute Physiology Score II; NA, not available.

**Table 4.** Patient characteristics (Study 2)

|  |  |  |  |  | Comorbidities |  |  |  |  |  |  |  |  |  | Gas6 |  |  | sAxl |  |  | Gas6-sAxl complexes | sMer |  | sTyro3 |  | SOFA | SIC | Outcome (Day 30) | Outcome (Day 90) |
|---|---|---|---|---|---|---|---|---|---|---|---|---|---|---|---|---|---|---|---|---|---|---|---|---|---|---|---|---|---|
| Patient No. | Age | Sex | Admission Source | Renal Failure | Myocardial Infarction | Cerebrovascular accident OR Transient ischemic attack | Congestive Heart Failure | Chronic Obstructive Pulmonary Disease | Liver disease | Diabetes mellitus | Leukemia | Primary site of infection | Hospital aquired infection | COVID-19 | ng/mL | % | nM | ng/mL | % | nM | nM | ng/mL | % | ng/mL | % | Score | Score |  |  |
| 1 | 60 | Male | IMC | Acute renal failure | No | Yes | No | No | Mild | End-organ damage | No | Unkown | No | No | 30.2 | 380 | 0.37 | 34.8 | 137 | 0.81 | 0.32 | 37.5 | 527 | 2.4 | 276 | 12 | 4 | Survival | Survival |
| 2 | 76 | Male | Other Hospital | Chronic renal failure | Yes | No | Yes | Yes | Mild | Diet-controlled | No | Unkown | No | No | 17.5 | 220 | 0.21 | 48.1 | 189 | 1.12 |  | 9.8 | 138 | 1.2 | 138 | 11 | 3 | Survival |  |
| 3 | 84 | Female | Emergency department | None | No | No | No | No | Mild | None | No | Gastrointestinal or biliary | No | No | 17.1 | 216 | 0.21 | 41.5 | 163 | 0.97 | 0.52 | 11 | 154 | 0.8 | 92 | 8 | 2 | Survival | Survival |
| 4 | 66 | Female | Emergency department | Acute renal failure | No | No | No | No | Mild | None | No | Respiratory | No | Yes | 38.5 | 485 | 0.47 | 28 | 110 | 0.65 | 0.27 | 20 | 281 | 1 | 115 | 12 | 4 | N/A | N/A |
| 5 | 42 | Male | Hospital ward | None | No | No | No | No | Mild | None | No | Gastrointestinal or biliary | Yes | No | 33.0 | 416 | 0.40 | 11.2 | 44 | 0.26 | 0.08 | 15.3 | 215 | 1.2 | 138 | 14 | 5 | Survival | Survival |
| 6 | 49 | Female | Emergency department | Acute on chronic renal failure | No | No | No | No | Mild | None | No | Urogenital | No | No | 26.6 | 335 | 0.32 | 32.5 | 128 | 0.76 | 0.57 | 25.9 | 364 | 1.5 | 172 | 13 | 4 | Survival | Survival |
| 7 | 66 | Female | Emergency department | Acute renal failure | No | No | No | No | Mild | None | Yes | Respiratory | No | No | 29.9 | 376 | 0.36 | 48.3 | 190 | 1.12 |  | 57 | 800 | 2.3 | 264 | 14 | 4 | Death |  |
| 8 | 79 | Male | Emergency department | None | No | No | No | Yes | Mild | Diet-controlled | No | Urogenital | No | No | 24.7 | 311 | 0.30 | 44.2 | 174 | 1.03 |  | 13.4 | 188 | 1 | 115 | 12 | 5 | Survival | Survival |
| 9 | 58 | Female | Emergency department | Acute renal failure | No | No | No | No | Mild | None | No | Respiratory | Yes | No |  |  | 0.25 |  |  | 0.47 | 0.07 |  |  |  |  |  |  | Survival | Survival |
| 10 | 63 | Male | Hospital ward | None | No | No | No | Yes | Mild | None | No | Respiratory | Yes | No | 16.3 | 206 | 0.20 | 27.1 | 107 | 0.63 | 0.44 | 6 | 84 | 1 | 115 | 11 | 2 | Survival | Survival |
| 11 | 77 | Female | Emergency department | None | No | No | No | No | Mild | None | No | Respiratory | No | No | 11.8 | 149 | 0.14 | 32.1 | 126 | 0.75 |  | 12.9 | 181 | 1.4 | 161 | 9 | 2 | Survival | Survival |
| 12 | 64 | Male | Emergency department | Unknown | No | No | No | Yes | Mild | None | No | Respiratory | No | No | 13.4 | 169 | 0.16 | 15.1 | 59 | 0.35 | 0.21 | 10.1 | 142 | 1.4 | 161 | 12 | 3 | Survival | Survival |
| 13 | 49 | Female | Hospital ward | Acute renal failure | No | No | No | No | Mild | None | No | Gastrointestinal or biliary | No | No | 44.4 | 559 | 0.54 | 21.9 | 86 | 0.51 | 0.26 | 25.2 | 354 | 1.8 | 207 | 16 | 4 | Survival | Survival |
| 14 | 70 | Female | Other Hospital | Acute renal failure | No | No | No | Yes | Mild | None | No | Prevertebral abscess DD intestinal necrosis | No | No | 21.2 | 267 | 0.26 | 46.1 | 181 | 1.07 | 0.24 | 28.4 | 399 | 1.8 | 207 | 17 | 4 | N/A | N/A |
| 15 | 75 | Female | Emergency department | Acute renal failure | No | Yes | No | No | Mild | Uncomplicated | No | Respiratory | No | No | 31.7 | 399 | 0.39 | 33.3 | 131 | 0.77 |  | 13.9 | 195 | 2.3 | 264 | 12 | 3 | Survival | Death |
| 16 | 29 | Male | Emergency department | Acute renal failure | No | No | No | No | Mild | None | No | Neuro | No | No | 14.5 | 183 | 0.18 | 27.5 | 108 | 0.64 | 0.36 | 11.6 | 163 | 1.4 | 161 | 12 | 4 | Survival | Survival |
| 17 | 67 | Male | Other Hospital | Acute renal failure | No | No | No | No | Mild | None | No | Respiratory | No | No | 20.9 | 263 | 0.25 | 25.1 | 99 | 0.58 |  | 11.8 | 166 | 1.7 | 195 | 10 | 2 | Survival | Survival |
| 18 | 66 | Female | Emergency department | None | No | No | No | Yes | Mild | None | No | Respiratory | No | No |  |  |  |  |  |  |  |  |  |  |  | 11 |  | Death |  |
| 19 | 72 | Male | Other Hospital | Acute on chronic renal failure | No | Yes | No | No | Mild | None | No | Gastrointestinal or biliary | No | No | 29.5 | 372 | 0.36 | 42.1 | 165 | 0.98 |  | 16 | 225 | 1.5 | 172 | 17 | 5 | Death |  |
| 20 | 71 | Female | Hospital ward | Acute renal failure | No | No | No | No | Mild | None | Yes | Respiratory | Yes | No | 34.7 | 437 | 0.42 | 57.9 | 228 | 1.35 | 0.59 | 44 | 618 | 5 | 575 | 15 | 6 | Death |  |
| 21 | 87 | Male | IMC | Acute renal failure | Yes | Yes | Yes | No | Mild | None | No | Unkown | Yes | No | 14.9 | 188 | 0.18 | 34.7 | 136 | 0.81 |  | 20 | 281 | 1 | 115 | 14 | 2 | Death |  |
| 22 | 70 | Female | Recovery room | Acute renal failure | No | No | No | No | Moderate to severe | None | No | Soft tissue or skin | No | No | 49.4 | 622 | 0.60 | 85.7 | 337 | 1.99 |  | 51.9 | 729 | 2.2 | 253 | 16 | 6 | Death |  |
| 23 | 70 | Male | Emergency department | Acute on chronic renal failure | No | No | Yes | Other pulmonary disease | Mild | Uncomplicated | No | Primary bacteremia | No | No | 17.2 | 217 | 0.21 | 38.8 | 153 | 0.90 |  | 28.7 | 403 | 1.3 | 149 | 11 | 6 | Survival | Survival |
| 24 | 78 | Male | Hospital ward | Acute renal failure | Yes | No | Yes | Yes | Mild | Uncomplicated | No | Gastrointestinal or biliary | No | No | 11.3 | 142 | 0.14 | 19.6 | 77 | 0.46 |  | 10 | 140 | 0.8 | 92 | 13 |  | N/A | N/A |

Abbreviations: COVID-19, coronavirus disease 2019; IMC, Intermediate Care; SIC, sepsis-induced coagulopathy; SOFA, Sequential Organ Failure Assessment.

To determine whether these observations could be reproduced in experimental models of infection, we used two murine models of microbial sepsis based on peritonitis. In the first model, *Gas6^+/+^* mice were inoculated intraperitoneally with the *E. coli* strain O18^57^. Plasma Gas6 levels increased by approximately 30% within 4 hours (n = 6) and remained elevated for at least 24 hours (**Fig. 1D**) In a second model that closely mimics human peritoneal sepsis—CLP^61,62^—plasma Gas6 concentrations were also significantly higher at 8 and 24 hours in mice with peritonitis compared with sham-operated controls (**Fig. 1E**). Owing to the severity of this model, longer-term follow-up of plasma Gas6 levels was not feasible. These data demonstrate that experimental microbial sepsis in mice recapitulates the sustained elevation of Gas6 observed in human sepsis.

We next assessed systemic inflammation in humans using the experimental endotoxemia model, the only controlled *in vivo* model available to study acute systemic inflammatory responses in healthy individuals^63^. Although this model has limitations—including the absence of capillary leakage, hemodynamic shock, and overt organ dysfunction, as well as a predominantly innate immune response with minimal adaptive contribution—it is highly standardized and reproducible. Healthy volunteers received an intravenous (i.v.) injection of LPS to induce a transient systemic inflammatory response. Plasma Gas6 levels were measured at baseline and every 30 minutes following LPS administration^58^. Gas6 concentrations increased rapidly after LPS injection, reaching a peak at approximately 90 minutes with an approximate twofold increase from baseline (P < 0.001, **Fig. 1F**), and remained elevated throughout the 360-minute observation period. These findings indicate that LPS directly triggers a rapid and sustained release of Gas6 into the circulation, consistent with the pattern observed in patients with sepsis.

We then applied an LPS-induced endotoxemia model in mice using doses of 25 or 50 mg/kg, resulting in approximately 25% (LD25) and 75% (LD75) mortality in *WT* mice, respectively. Under steady-state conditions, circulating Gas6 levels in *Gas6^+/+^* mice were 24 ± 2 ng/mL (= 0.33 ± 0.03 nM). Following LPS challenge, Gas6 concentrations increased rapidly, peaking at 1 hour to 55 ± 10 ng/mL (= 0.75 ± 0.37 nM) and 66 ± 6 ng/mL (=0.9 ± 0.08 nM) after 25 and 50 mg/kg LPS, respectively (**Fig. 1G**). Gas6 levels remained significantly elevated for up to 4 hours after injection (P < 0.0001 vs. baseline), followed by a gradual decline toward baseline values (**Fig. G**). Importantly, Gas6 concentrations were significantly higher after 50 mg/kg than after 25 mg/kg LPS (P < 0.01; **Fig. 1G**), indicating a dose-dependent response. Compared with humans, however, the decline in circulating Gas6 levels occurred more rapidly in mice.

Because protein S shares structural homology with Gas6 and can also bind TAM receptors, we measured plasma protein S levels in parallel. In contrast to the rapid increase in Gas6 following LPS administration, protein S levels showed a slight early decrease and remained below baseline for at least 12 hours (1 hour vs. 4 hours, P < 0.05; **Fig. 1H**). Thus, exposure to a bacterial product induces a prompt and robust increase in circulating Gas6, whereas protein S does not exhibit a similar response.

Taken together, these findings demonstrate that Gas6 is rapidly and robustly induced during systemic inflammation in both humans and mice. Plasma Gas6 levels rise early in sepsis and experimental endotoxemia, correlate with disease severity, and are associated with increased mortality in patients. The conserved and dose-dependent nature of this response across species supports a pathophysiological role for Gas6 in sepsis and highlights its potential as a biomarker of disease severity and outcome.

### Genetic loss of Gas6/TAM heightens endotoxemia and infection susceptibility whereas exogenous Gas6 confers protection

Since only Gas6, and not protein S, was increased in sepsis, we examined whether endogenous Gas6 is an active player in the endotoxic response by using *Gas6^+/+^*, *Gas6^+/-^* and *Gas6^-/-^* mice models. After LPS injection, *Gas6^-/-^* mice showed an increased vulnerability compared to both *Gas6^+/+^* and *Gas6^+/-^* mice (**Fig. 2A-B**). In particular, whereas most of the *Gas6^-/-^* mice succumbed in the first 48 hours after 25 mg/kg LPS treatment, 73% of the *Gas6^+/+^* mice survived (**Fig. 2A**). Using higher LPS dosage (50 mg/kg), all *Gas6^-/-^* mice died within 48 hours (**Fig. 2B**) whereas 25% of *Gas6^+/+^* mice were still alive; the median survival was 34 hours for *Gas6^+/+^* versus 14.5 hours for *Gas6^-/-^* mice. Survival after LPS challenge in *Gas6^+/-^* mice, whose circulating Gas6 was about 50% of the *Gas6^+/+^* level (**Fig. 2C**) did not differ from *Gas6^+/+^*. In the CLP model of peritonitis^61,62^, *Gas6^-/-^* mice were more susceptible than *Gas6^+/+^* mice: 83% of *Gas6^-/-^* mice died within 48 hours of surgery, compared with 55% of *Gas6^+/+^* mice (**Fig. 2D**), while all sham-operated mice survived. These findings indicate that Gas6 deficiency increases vulnerability to endotoxemia and microbial sepsis.

**FIGURE 2:**
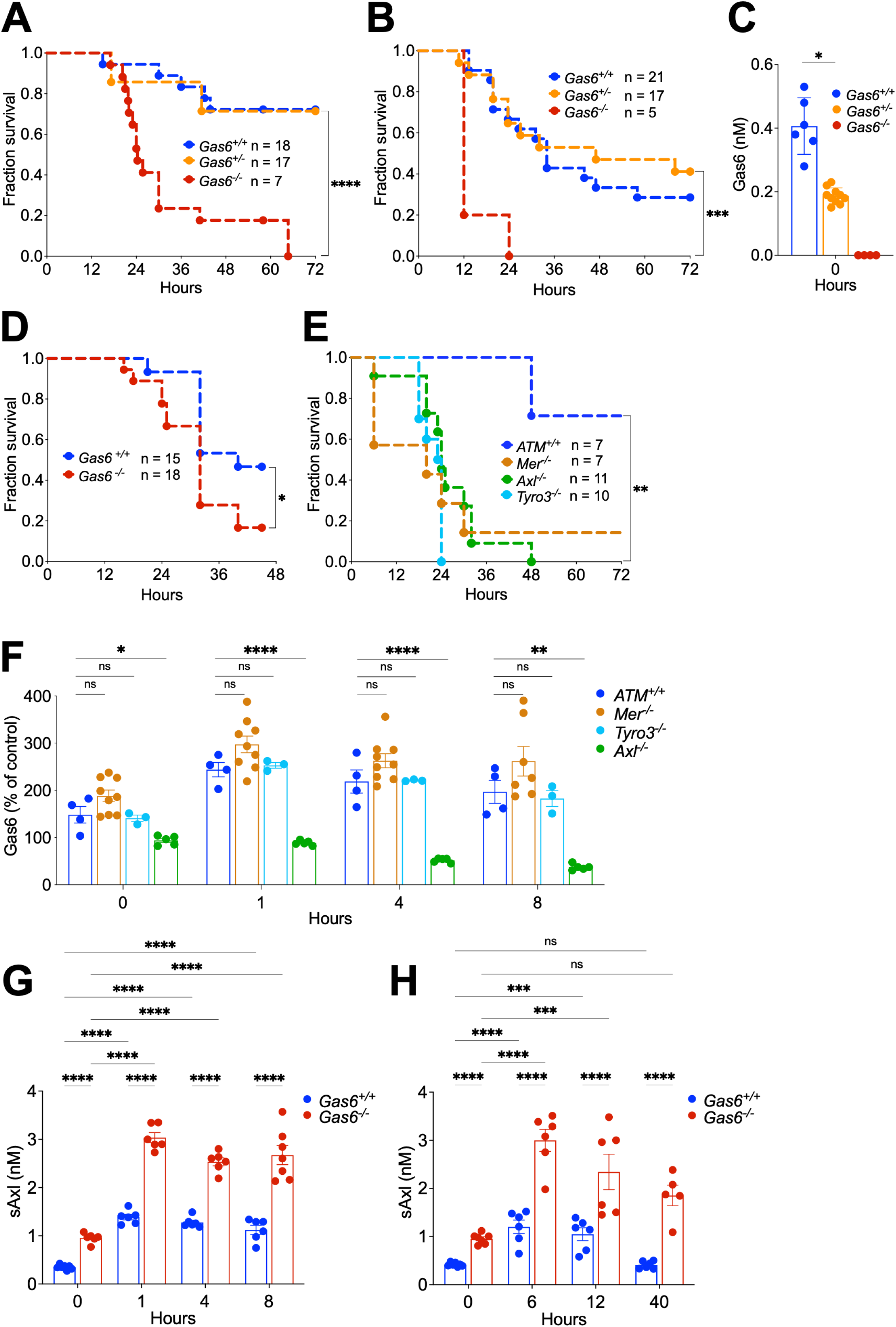
Genetic loss of Gas6/TAM heightens endotoxemia and infection susceptibility whereas exogenous Gas6 confers protection. **(A-B)** Kaplan-Meier plots of the survival of *Gas6^+/+^*, *Gas6^+/-^* and *Gas6^-/-^* mice treated with LPS 25 mg/kg **(A)** and 50 mg/kg **(B)** intraperitoneally (i.p.). Data are pooled from multiple independent experiments. Log -rank Mantel-Cox test. **(C)**, Gas6 circulating level in *Gas6^+/+^*, *Gas6^+/-^* and *Gas6^-/-^* mice, mean ± SEM, Student *t* test. **(D)**, CLP survival in *Gas6^-/-^* and *Gas6^+/+^* mice. Data from three independent experiments. Log -rank Mantel-Cox test. **(E)**, Kaplan-Meier plots of the survival of wild-type (*ATM^+/+^*) and mice lacking any one of the cognate Gas6 receptors (*Axl^-/-^*, *Mer^-/-^* and *Tyro3^-/-^* mice) treated with 25 mg/kg LPS. Data from two independent experiments. Log -rank Mantel-Cox test. **(F)**, Gas6 circulating level in *ATM^+/+^* and *Axl^-/-^*, *Mer^-/-^* and *Tyro3^-/-^* mice mice in steady-state (0 hours) and at 1-8 hours after LPS challenge (25 mg/kg). Dunnett’s nultiple comparison test. **(G-H)** sAxl plasma levels in *Gas6^+/+^* and *Gas6^-/-^* at baseline and after LPS challenge: 50 mg/kg **(G)**, and 10 mg/kg **(H)**. Tukey’s multiple comparison test. **(I-J),** sMer plasma levels in *Gas6^+/+^* and *Gas6^-/-^* at baseline and after LPS challenge: 50 mg/kg **(I)**, and 10 mg/kg **(J)**. Tukey’s multiple comparison test. **(K)**, Urine levels of Gas6 in *Gas6^+/+^* mice at baseline and after LPS 50 mg/kg. Dunnett’s multiple comparison test. **(L),** Urine levels of sAxl in *Gas6^+/+^* and *Gas6^-/-^* mice at baseline and after LPS. Tukey’s multiple comparison test **(M)**, Urine levels of sMer in *Gas6^+/+^* and *Gas6^-/-^* mice at baseline and after LPS. Tukey’s multiple comparison test. **(N)**, Plasma levels of Gas6 at baseline and after LPS challenge (10 mg/kg) with or without rGas6 (10 µg i.v.). Tukey’s multiple comparison test. **(O)**, Kaplan-Meier plots of the survival of *Gas6^+/+^* treated with LPS 10 mg/kg without or with rGas6 (10 µg i.v.) administered at different timepoint following LPS injection. Log -rank Mantel-Cox test. **(P)**, Plasma levels of sAxl at baseline and 4 hours after LPS challenge (10 mg/kg) with or without rGas6 (10 µg i.v.). Tukey’s multiple comparison test. **(Q)**, Plasma levels of Gas6, soluble Gas6–sAxl complexes, and sAxl in patients with septic shock. Tukey’s multiple comparison test. ns, non-significant, \**P* < 0.05, \*\**P* < 0.01, \*\*\**P* < 0.001,\*\*\*\**P* < 0.0001

To further define the role of Gas6 and its signaling through TAM receptors in the innate response to endotoxemia, we challenged mice lacking individual TAM receptors with LPS 25 mg/kg (**Fig. 2E**). *Axl^-/-^ Tyro3^-/-^*, or *Mer^-/-^* mice were all significantly more susceptible to LPS 25 mg/kg than wild-type (*ATM^+/+^*) controls (n = 7–11; all P < 0.01), with no differences in sensitivity among the knockout strains (P > 0.05). The findings in *Mer^-/-^* mice are consistent with previous reports by Camenisch et al.^49^. At baseline, *Axl^-/-^* but not *Mer^-/-^* and *Tyro3^-/-^* mice had lower plasma Gas6 levels than *ATM^+/+^* mice (**Fig. 2F**). Following LPS administration (25 mg/kg), Gas6 levels increased in *ATM^+/+^*, *Mer^-/-^* and *Tyro3^-/-^* mice and remained elevated for the next 8 hours. In *Axl^-/-^* mice, Gas6 plasma levels did not raise at one hour and then, declined progressively during the first 8 hours (Timepoint 0 *versus* 4 hours: P < 0.0001; 0 *versus* 8 hours: P < 0.0001; 1 *versus* 4 hours: P < 0.0001; 1 *versus* 8 hours: P < 0.0001; 4 versus 8 hours: P < 0.05, Tukey’s multiple comparisons test). These results suggest that the systemic rise in Gas6 during inflammation depends on the presence of Axl.

During inflammation, TAM receptors can be cleaved and released into the circulation. At baseline, *Gas6^-/-^* mice displayed higher soluble Axl (sAxl) levels than *Gas6^+/+^* mice (**Fig. 2G**). Following administration of 50 mg/kg LPS, sAxl levels increased in both genotypes during the first 7 hours and were approximately twofold higher in *Gas6^-/-^* mice. After a lower LPS dose (10 mg/kg) allowing a prolonged survival, sAxl was monitored for up to 40 hours (**Fig. 2H**). Although levels gradually declined over time both in *Gas6^+/+^* and *Gas6^-/-^* mice, they remained elevated in *Gas6^+/+^* mice before returning to baseline by 40 hours, whereas they remained elevated in *Gas6^-/-^* mice. In contrast, soluble Mer (sMer) levels did not change over time neither after the 50 mg/kg nor the 10 mg/kg LPS challenges (**Fig. 2I-J**), and soluble Tyro3 could not be detected in murine plasma by ELISA. Taken together, these data indicate a tight relationship between circulating Gas6 and sAxl. Consistent with this, reduced Gas6 levels were observed in *Axl^-/-^* mice at baseline and during LPS challenge, whereas sAxl levels were increased in the absence of Gas6, supporting an interdependent regulation of these two molecules.

To investigate the mechanisms underlying Gas6, sAxl, and sMer elimination during endotoxemia, we quantified their urinary concentrations at multiple time points following LPS administration. At baseline, urinary Gas6 levels were comparable to those in plasma. However, in contrast to the marked changes observed in circulating Gas6 during LPS challenge, urinary Gas6 concentrations remained stable (**Fig. 2K**). This dissociation suggests that urinary Gas6 is largely derived from constitutive local secretion within the urinary tract rather than from glomerular filtration of circulating Gas6, and that this local production is not significantly modulated by systemic inflammation.

Baseline urinary sAxl levels were similar to plasma concentrations, consistent with partial renal handling under steady-state conditions. Following administration of 50 mg/kg LPS, urinary sAxl increased markedly, particularly in *Gas6^-/-^* mice (**Fig. 2L**). This transient rise supports enhanced renal clearance as a major pathway for sAxl elimination during acute inflammation, potentially driven by increased circulating levels and/or altered glomerular permeability. Notably, urinary sAxl concentrations returned to baseline by 4 hours, indicating rapid normalization of renal handling mechanisms.

In *Gas6^-/-^* mice, urinary sAxl levels matched plasma concentrations at baseline, exceeded plasma levels approximately twofold at 1 hour, and subsequently realigned with plasma concentrations at 4 and 8 hours (**Fig. 2G, K**). Together, these data support a dynamically regulated, inflammation-dependent renal clearance mechanism for sAxl that is accentuated in the absence of Gas6.

This pronounced rise indicates that excess circulating sAxl, as observed during endotoxemia, can be effectively cleared by the kidneys. Urinary sMer levels also increased after LPS exposure but to a lesser extent (**Fig. 2M**), indicating a more modest degree of renal elimination compared with sAxl.

Because plasma Gas6 levels are elevated in septic patients who do not survive compared with survivors,^39^ we investigated the effect of exogenous Gas6 administration in *Gas6^+/+^* mice challenged with LPS. Recombinant murine Gas6 (rGas6) was administered 4 hours after LPS injection at a dose sufficient to almost doubling circulating Gas6 concentration (**Fig. 2N**). Remarkably, all mice treated with rGas6 survived the first 24 hours following LPS administration (10 mg/kg), indicating a protective effect of high-dose Gas6 (**Fig. 2O**). Notably, a trend toward lower sAxl levels was observed 4 hours after LPS injection in mice treated with a high-dose of rGas6 (**Fig. 2P**).

These findings suggest that although Gas6 levels rise during systemic inflammation such as sepsis, this increase is not inherently deleterious. Rather, elevated Gas6 may represent a compensatory component of the inflammatory response, arguing against the notion that Gas6 itself is a pathogenic mediator or a suitable target for therapeutic inhibition in acute systemic inflammation.

To better understand the circulating equilibrium between Gas6 and sAxl, we quantified both proteins in plasma of *Gas6^+/+^* mice at steady-state, after LPS administration, and following rGas6 administration during endotoxemia (**Fig. 2N and P**). At steady-state, circulating Gas6 concentrations averaged 29.7 ng/mL, corresponding to approximately 0.40 nM assuming a molecular weight of 73 kDa. In parallel, sAxl levels averaged 28.3 ng/mL, corresponding to 0.36 nM assuming a molecular weight of 70 kDa. Thus, under physiological conditions, Gas6 and sAxl were present at nearly equimolar concentrations (Gas6/sAxl ≈ 1.11). Assuming a high-affinity 1:1 interaction between Gas6 and sAxl, as previously proposed for circulating Gas6–Axl complexes^64^, the maximal concentration of soluble complexes would be limited by the lower component (0.36 nM). Based on this estimate, approximately 0.038 nM Gas6 would remain unbound, suggesting that about 90% of circulating Gas6 could theoretically be complexed with sAxl under baseline conditions.

Following LPS administration, plasma Gas6 increased to 68 ng/mL (0.907 nM, **Fig. 2N**), while sAxl increased in parallel to 68.2 ng/mL (0.863 nM, **Fig. 2P**). Despite this marked induction of both molecules, their molar ratio remained close to unity (Gas6/sAxl ≈ 1.05). Under these conditions, the maximal concentration of soluble Gas6–sAxl complexes would be predicted to reach 0.863 nM, leaving approximately 0.044 nM free Gas6 in circulation. Thus, although endotoxemia substantially increases circulating Gas6 levels, the concomitant rise in sAxl maintains a near-stoichiometric equilibrium in which the vast majority of Gas6 could theoretically sequestered within the soluble Gas6–Axl complexes.

In contrast, administration of rGas6 markedly altered this equilibrium. Thirty minutes after rGas6 injection, measurable sAxl levels decreased to 45.5 ng/mL (0.576 nM, **Fig. 2P**), representing a 33% reduction compared with mice challenged with LPS alone (0.863 nM). Because total Gas6 concentrations following rGas6 administration necessarily exceed the endotoxemia level of 0.907 nM, the maximal concentration of soluble Gas6–sAxl complexes becomes limited by the reduced sAxl pool (0.576 nM). Even under the conservative assumption that total Gas6 remained at 0.907 nM, the minimal concentration of unbound Gas6 would therefore increase to >0.331 nM, representing at least a 7.5-fold increase in free ligand availability compared with endotoxemia alone (0.331 nM vs. 0.044 nM). These calculations suggest that rGas6 administration exceeds the buffering capacity of the sAxl compartment, shifting the circulating system from a tightly buffered state toward a ligand-excess state. This shift would be expected to increase the fraction of Gas6 available to engage membrane TAM receptors, providing a plausible mechanistic explanation for the protective effects of rGas6 observed during endotoxemia.

We next measured plasma Gas6, sAxl and soluble Gas6-sAxl complexes in healthy volunteers and in patients with septic shock from Study 2 (**Fig. 2Q**). In healthy volunteers (n = 4), the concentrations of Gas6, sAxl and soluble Gas6-sAxl complexes were comparable (Gas6 vs sAxl, Gas6 vs Gas6-sAxl, sAxl vs Gas6-sAxl, Turkey’s multiple comparison test, P = ns). These results are consistent with the mouse data and support the notion that circulating Gas6 is largely bound to sAxl under physiological conditions.

In patients with septic shock (n = 12), plasma Gas6 concentrations were higher than in healthy volunteers, as expected. Gas6 and Gas6–sAxl complex concentrations were similar, while sAxl levels showed a slight excess relative to Gas6–sAxl complexes. These findings suggest that circulating Gas6 does not fully saturate the available sAxl pool, leaving little or no free Gas6 available to engage cellular TAM receptors.

Overall, deficiency of Gas6 or any TAM receptor increased susceptibility to endotoxemia, and Gas6 deficiency additionally heightened vulnerability to infection. These results suggest a functional interplay between Gas6 and sAxl in regulating TAM receptor signaling. Administration of high-dose Gas6 appears to overcome the buffering capacity of soluble Axl, thereby increasing the fraction of bioavailable Gas6 capable of activating membrane TAM receptors.

### *Gas6^-/-^* mice exhibit exacerbated hypotension and cytokine responses during endotoxemia

In rodents, LPS promotes hypotension accompanied by the release of pro-inflammatory cytokines and nitric oxide (NO), and by the induction of nitric oxide synthase expression. These events finally result in the onset of acute pulmonary edema, myocardial dysfunction and death^65,66^. To investigate whether *Gas6^-/-^* mice were more susceptible to hypotension during endotoxemia, arterial blood pressure was measured. *Gas6^-/-^* and *Gas6^+/+^* mice did not differ at steady state; however, 6 hours after LPS injection, mean blood pressure was significantly reduced in *Gas6^-/-^* mice (**Fig. 3A-C**).

**FIGURE 3:**
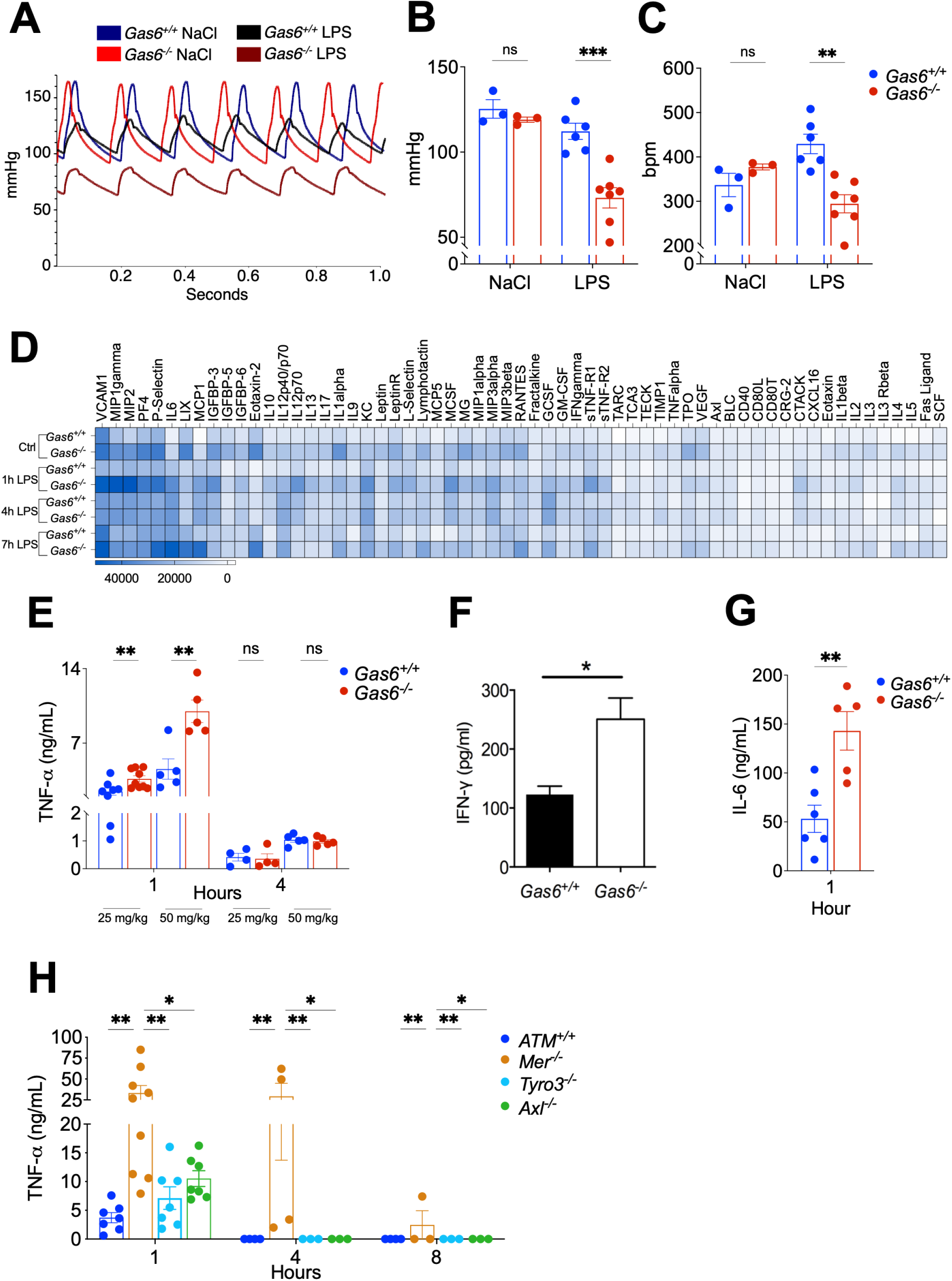
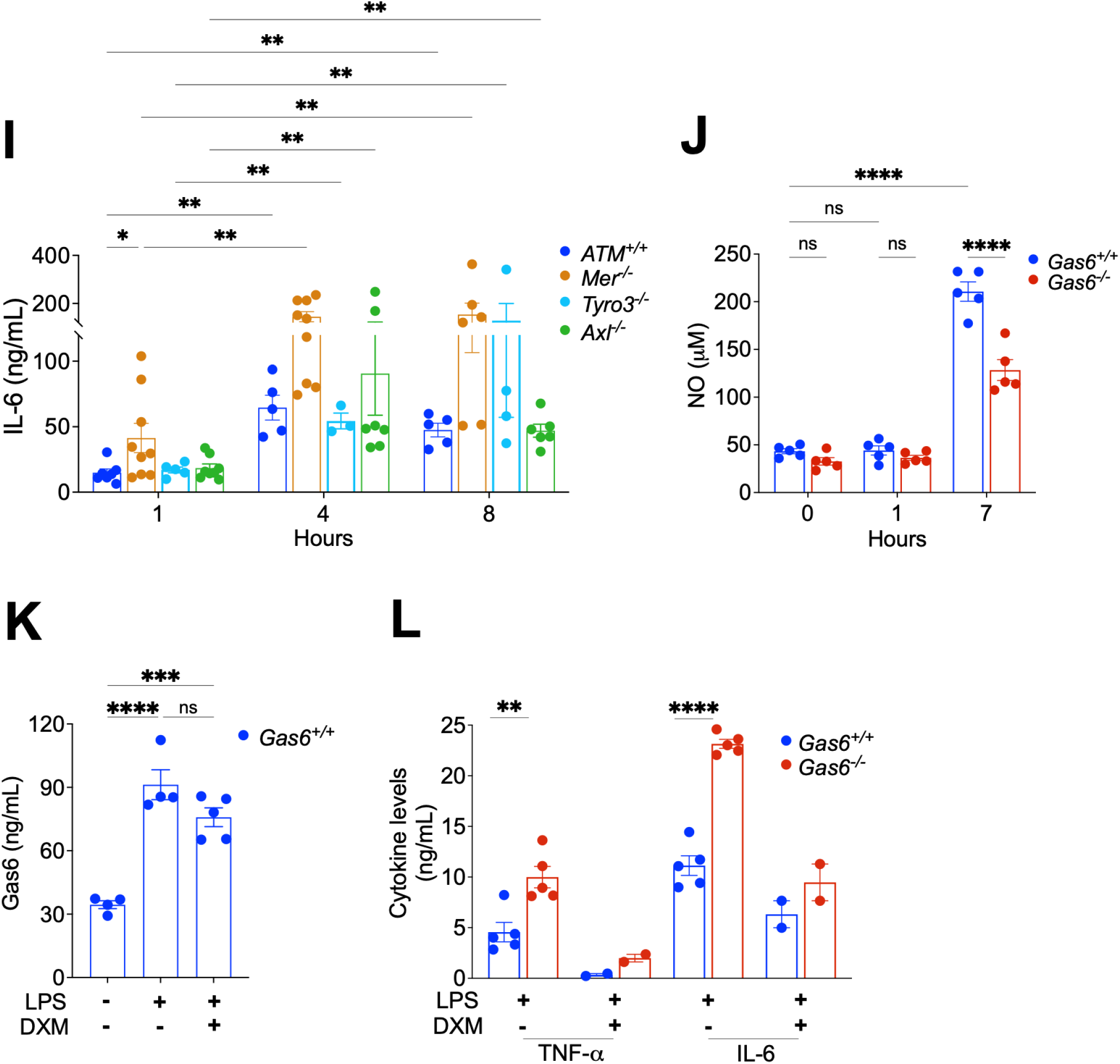
*Gas6^-/-^* mice exhibit exacerbated hypotension and cytokine responses during endotoxemia. **(A)**, Representative traces of arterial blood pressure measured in mice 6 hours after i.p. injection of NaCl 0.9% or 40 mg/kg LPS. Data from two independent experiments. **(B-C)**, Mean blood pressure **(B)** measured in the carotid artery and heart rate **(C)** did not differ between *Gas6^+/+^* (n = 3) and *Gas6^-/-^* mice (n = 3) in steady state (mean ± SEM) but was lower in *Gas6^-/-^* mice (*n* = 7) compared to *Gas6^+/+^* mice (*n* = 6) 6 hours after LPS injection (40 mg/kg). Data from two independent experiments. Šídák’s multiple comparisons test. **(D)**, Cytokines antibody array was performed using pool of *Gas6^-/-^* and *Gas6^+/+^* plasma (n=8 mice each group) 1 hour after LPS injection (50 mg/kg). The relative expression of the 62 cytokines has been assessed in arbitrary units (AU) and show in a heat map. Details about the cytokines are listed in **Table 1**. (**E**), Plasmatic TNF-α in *Gas6^-/-^* and *Gas6^+/+^* mice 1 and 4 hours after LPS 25 mg/kg. Data from two independent experiments. Tukey’s multiple comparison test. (**F**), IFN-γ levels in *Gas6^-/-^* and *Gas6^+/+^* mice 1 hour after LPS injection. Data from two independent experiments. (**G**), IL-6 levels measured in the peritonitis model triggered by E. coli inocula, higher dose of E. coli ( 2.5*106 CFU) in *Gas6^-/-^* than *Gas6^+/+^* mice, n = 5-6. Unpaired test. (**H-I**), Plasmatic TNF-α and IL-6 in *ATM^+/+^*, *Axl^-/-^*, *Mer^-/-^*, *Tyro3^-/-^* mice 1, 4 and 8 hours after LPS injection (25 mg/kg). Tukey’s multiple comparisons test. (**J**), NO concentration in the blood of *Gas6^+/+^* and *Gas6^-/-^*, at different time points after LPS injection (25 mg/kg). Šídák’s multiple comparisons test. (**K**) Dexamethasone (DXM) does not control plasmatic Gas6. (**L**) The absence of Gas6 does not impair the immunomodulatory effect of DXM (LPS 50 mg/kg, 1 hour). ns, non significant, \**P* < 0.05, \*\**P* < 0.01, \*\*\**P* < 0.001,\*\*\*\**P* < 0.0001

Sixty-two pro-inflammatory and immunomodulatory cytokines were quantified in plasma in a relative scale manner at different time points after LPS injection (50 mg/kg i.p.) using an antibody array. Cytokines levels were generally higher in *Gas6^-/-^* mice 1 hour after LPS challenge (**Fig. 3D**), several cytokines being at least 50% higher in *Gas6^-/-^* than *Gas6^+/+^* plasma after LPS injection (after 1 hour: 46/62 cytokines, 4 hours: 6/62, 7 hours: 29/62, **Table 5**) but also at steady-state (52/62)^67^. Indeed, 1 hour after LPS injection, IFN-γ, IL-10, IL-12p40/p70, IL-12 p70, IL-1β, IL-6, KC were at least 50% higher in *Gas6^-/-^* than *Gas6^+/+^* plasma. After 4 hours, IL-1β and TNF-α, and after 7 hours, IL-10, MCP1 and TARC were at least 50% higher. Antigenic assays for TNF-α and IFN-γ further confirmed the increased concentration of both cytokines in *Gas6^-/-^* 1 hour after LPS injection (**Fig. 3E-F**). In the peritonitis model triggered by *E. coli* inocula, higher dose of *E. coli* (2.5*10^6^ CFU) induced much higher levels of IL-6 in *Gas6^-/-^* than *Gas6^+/+^* mice (143 ± 20 versus 53 ± 14 ng/mL, *P* = .004, *n* = 5-6) (**Fig. 3G**). In addition, we measured TNF-α levels in *Axl^-/-^*, *Tyro3^-/-^*, and *Mer^-/-^* mice following LPS challenge (25 mg/kg). Both *Axl^-/-^* mice and *Mer^-/-^* mice mice showed 3 to 10-fold increase of TNF-α levels compared to *ATM^+/+^* controls, whereas TNF-α levels in *Tyro3^-/-^* mice were comparable to those observed in *ATM^+/+^* mice (**Fig. 3H**). Similarly, IL-6 levels increased significantly at 1 hour in *Mer^-/-^* mice and rose significantly in *ATM^+/+^, Axl^-/-^*, *Tyro3^-/-^*, and *Mer^-/-^* mice at 4 and 8 hours after LPS administration. Notably, the increase in IL-6 was markedly greater in *Mer^-/-^* mice than in the other genotypes. Together, these findings indicate that among the TAM receptor knockouts, *Mer^−/−^* mice display the most rapid and pronounced inflammatory response to LPS, highlighting the critical role of MerTK in regulating acute systemic inflammation.

**Table 5.**
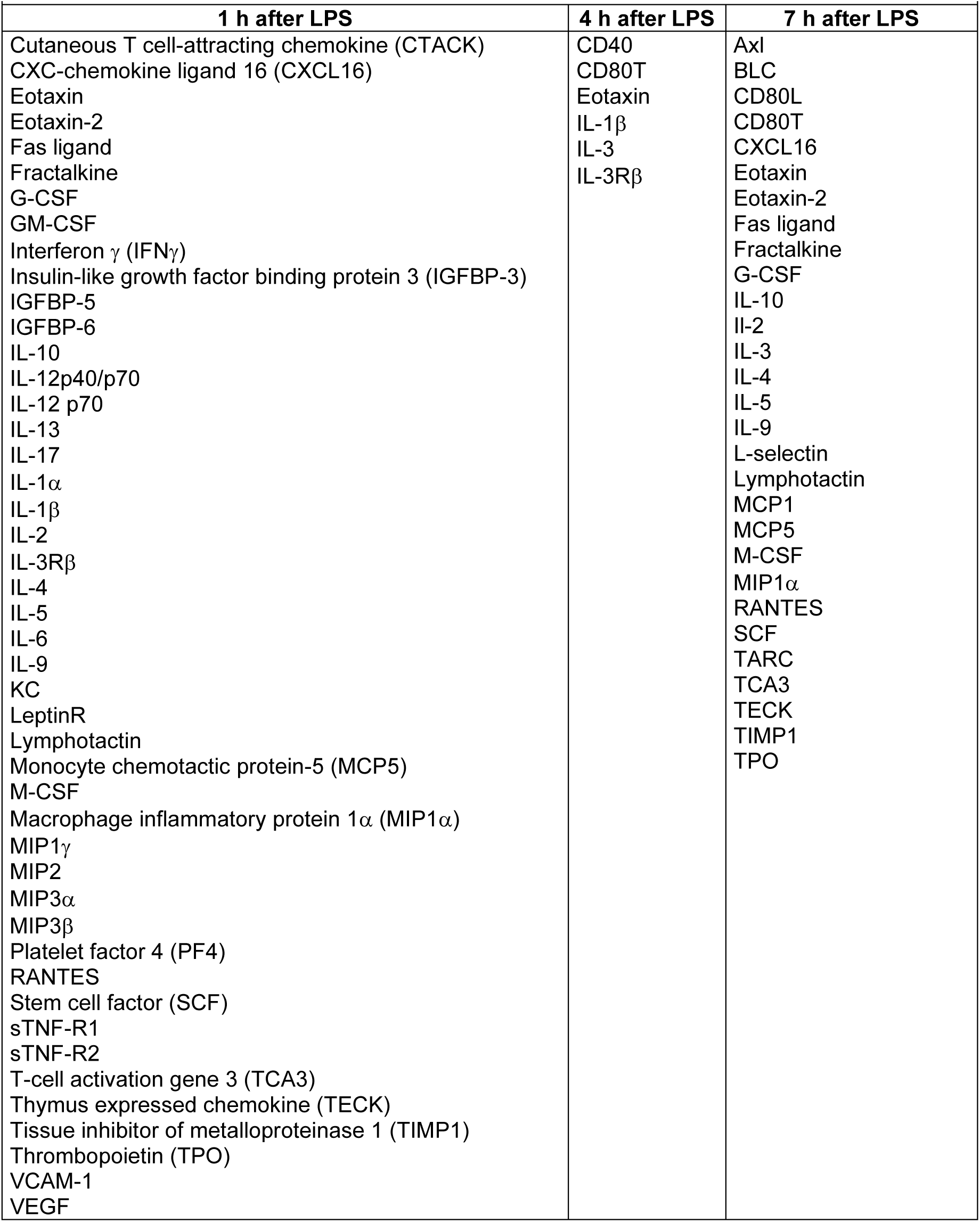
Cytokines at least 50% higher in plasma of *Gas6^-/-^* than *Gas6^+/+^* mice after LPS injection (50 mg/kg, intraperitoneally)

| 1 h after LPS | 4 h after LPS | 7 h after LPS |
|---|---|---|
| Cutaneous T cell-attracting chemokine (CTACK) | CD40 | Axl |
| CXC-chemokine ligand 16 (CXCL16) | CD80T | BLC |
| Eotaxin | Eotaxin | CD80L |
| Eotaxin-2 | IL-1β | CD80T |
| Fas ligand | IL-3 | CXCL16 |
| Fractalkine | IL-3Rβ | Eotaxin |
| G-CSF |  | Eotaxin-2 |
| GM-CSF |  | Fas ligand |
| Interferon γ (IFNγ) |  | Fractalkine |
| Insulin-like growth factor binding protein 3 (IGFBP-3) |  | G-CSF |
| IGFBP-5 |  | IL-10 |
| IGFBP-6 |  | Il-2 |
| IL-10 |  | IL-3 |
| IL-12p40/p70 |  | IL-4 |
| IL-12 p70 |  | IL-5 |
| IL-13 |  | IL-9 |
| IL-17 |  | L-selectin |
| IL-1α |  | Lymphotactin |
| IL-1β |  | MCP1 |
| IL-2 |  | MCP5 |
| IL-3Rβ |  | M-CSF |
| IL-4 |  | MIP1α |
| IL-5 |  | RANTES |
| IL-6 |  | SCF |
| IL-9 |  | TARC |
| KC |  | TCA3 |
| LeptinR |  | TECK |
| Lymphotactin |  | TIMP1 |
| Monocyte chemotactic protein-5 (MCP5) |  | TPO |
| M-CSF |  |  |
| Macrophage inflammatory protein 1α (MIP1α) |  |  |
| MIP1γ |  |  |
| MIP2 |  |  |
| MIP3α |  |  |
| MIP3β |  |  |
| Platelet factor 4 (PF4) |  |  |
| RANTES |  |  |
| Stem cell factor (SCF) |  |  |
| sTNF-R1 |  |  |
| sTNF-R2 |  |  |
| T-cell activation gene 3 (TCA3) |  |  |
| Thymus expressed chemokine (TECK) |  |  |
| Tissue inhibitor of metalloproteinase 1 (TIMP1) |  |  |
| Thrombopoietin (TPO) |  |  |
| VCAM-1 |  |  |
| VEGF |  |  |

Although NO enhances bacterial destruction, it is also well known to promote the vasodilation. We therefore evaluated the total amount of NO by measuring nitrate and nitrite concentrations in plasma. In WT mice, NO increased drastically after LPS injection (7 hours, **Fig. 3J**). In comparison, NO level increased only slightly in *Gas6^-/-^* mice.

Since steroids inhibit inflammatory cytokines, we assessed whether dexamethasone (DXM) affects Gas6 concentration in murine plasma or whether the lack of Gas6 would impair the immunomodulatory effect of the glucocorticoid. Intraperitoneal DXM injection in the presence or absence of LPS did not modify plasma Gas6 in comparison to vehicle controls (**Fig. 3K**). Moreover, in the absence of Gas6, DXM was still a potent inflammatory cytokines inhibitor (**Fig. 3L**).

Thus, during endotoxemia, numerous circulating pro- and anti-inflammatory cytokines were higher in *Gas6^-/-^* than *Gas6^+/+^* mice. In addition, TNF-α was higher in the absence of *Axl* or *Mer*, although the absence of *Tyro3* appeared to have less significance on TNF-α levels. However, NO levels were higher in *Gas6^+/+^* than *Gas6^-/-^* mice. DXM treatment, however, did not modify the release of Gas6 in plasma upon LPS challenge.

### Monocytes/macrophages lacking *Gas6* are activated and overproduce cytokines during endotoxemia

Although total peripheral blood leukocyte count and lymphocyte count were comparable in *Gas6^+/+^* and *Gas6^-/-^* 24 hours after LPS administration (50 mg/kg), the number of circulating monocytes was significantly higher in *Gas6^-/-^* than in *Gas6^+/+^* mice (**Fig. 4A**). To further characterize leukocyte population during endotoxemia, we performed mass cytometry analysis. Data were analyzed using unsupervised dimensionality reduction (t-SNE), which enables discrimination of major immune cell populations based on their phenotypic expression profiles (**Fig. 4B**). This analysis revealed that the absolute numbers of monocytes, neutrophils, NK cells, B cells, T helper cells, and cytotoxic T cells were comparable between *Gas6^+/+^* and *Gas6^−/−^* mice at steady-state and at 1.5 and 4 hours after LPS administration (50 mg/kg) (**Fig. 4C**). Following LPS challenge, monocyte counts decreased at 1.5 hours compared with steady state in both genotypes. In *Gas6^+/+^* mice, monocyte numbers returned to baseline by 4 hours, whereas in *Gas6^−/−^* mice they remained below steady-state levels. In contrast to monocytes, neutrophils increased more rapidly in *Gas6^−/−^* mice than in *Gas6^+/+^* mice, with significantly higher counts already observed at 1.5 hours. By 4 hours post-LPS, neutrophil numbers were markedly elevated relative to baseline in both genotypes and reached comparable levels. NK cell numbers increased in both genotypes at 4 hours compared with steady state. B cell counts progressively declined during the LPS challenge, with a similar decrease observed in both genotypes up to 4 hours. T helper cell numbers increased at 1.5 hours in Gas6+/+ mice relative to steady state but did not significantly change in *Gas6^−/−^* mice. At 4 hours, T helper cell counts were lower than both baseline and 1.5-hour levels in both genotypes. Similarly, cytotoxic T cells increased at 1.5 hours in *Gas6^+/+^* mice but not in *Gas6^−/−^* mice; however, cell counts in both genotypes returned to baseline by 4 hours.

**FIGURE 4:**
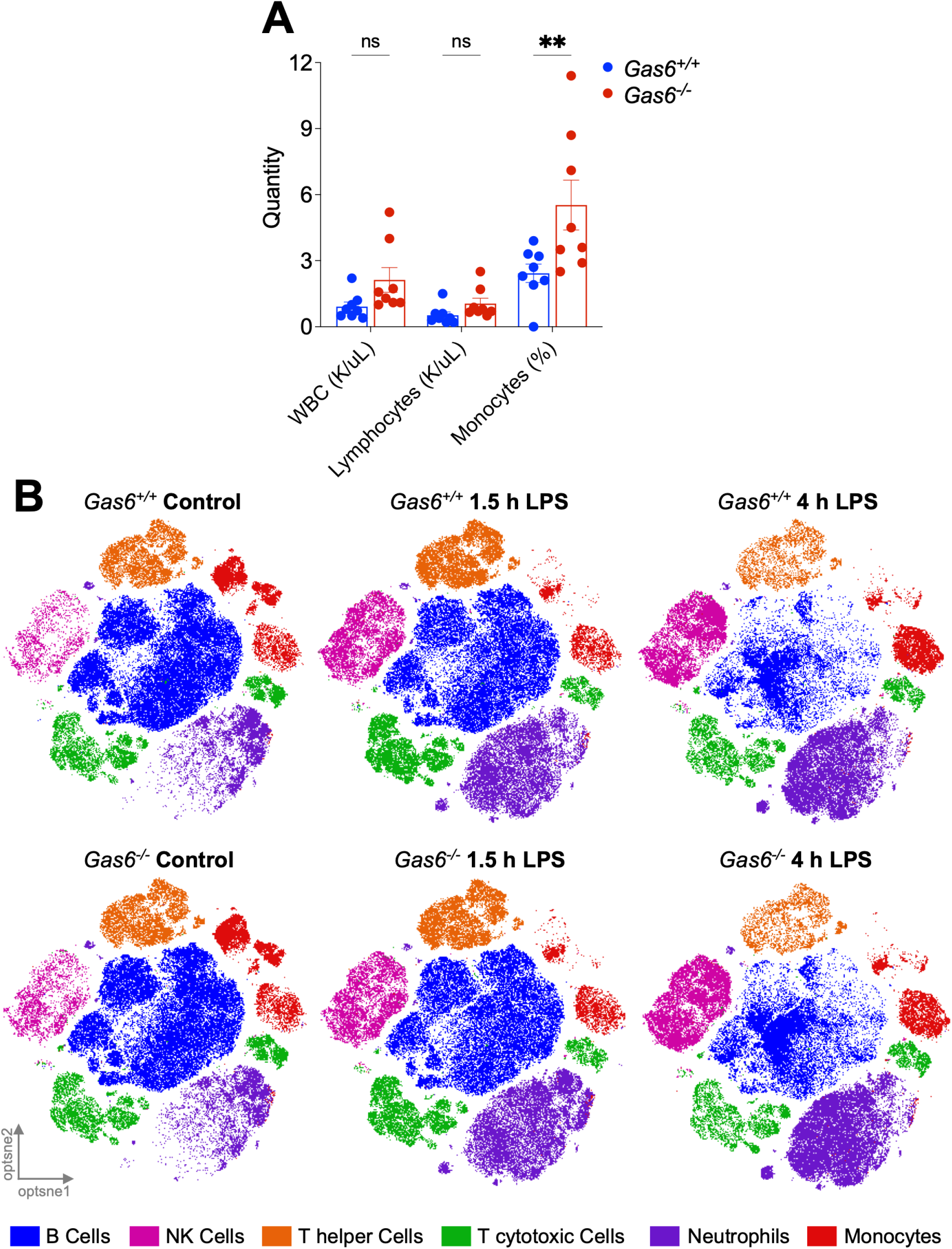
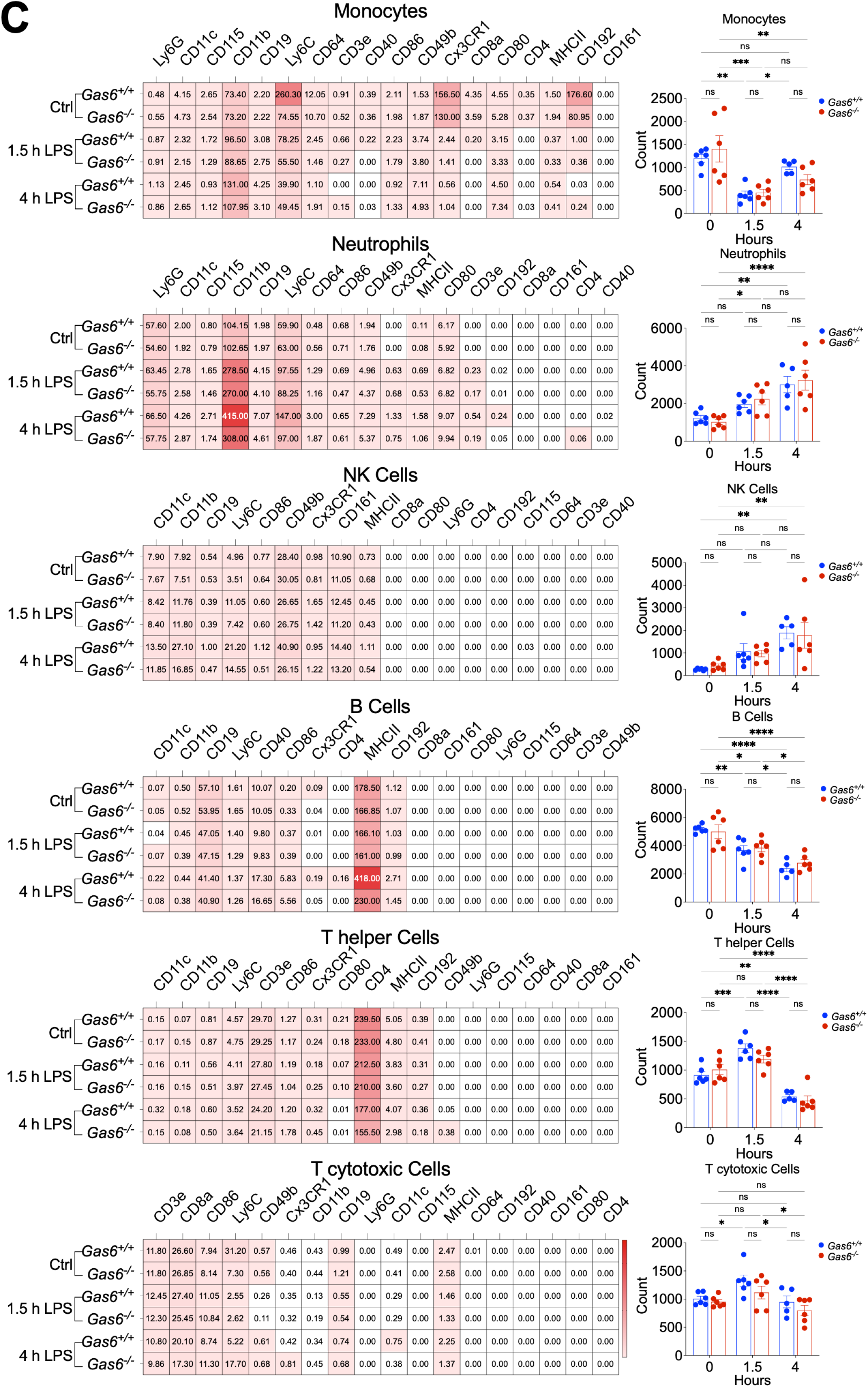
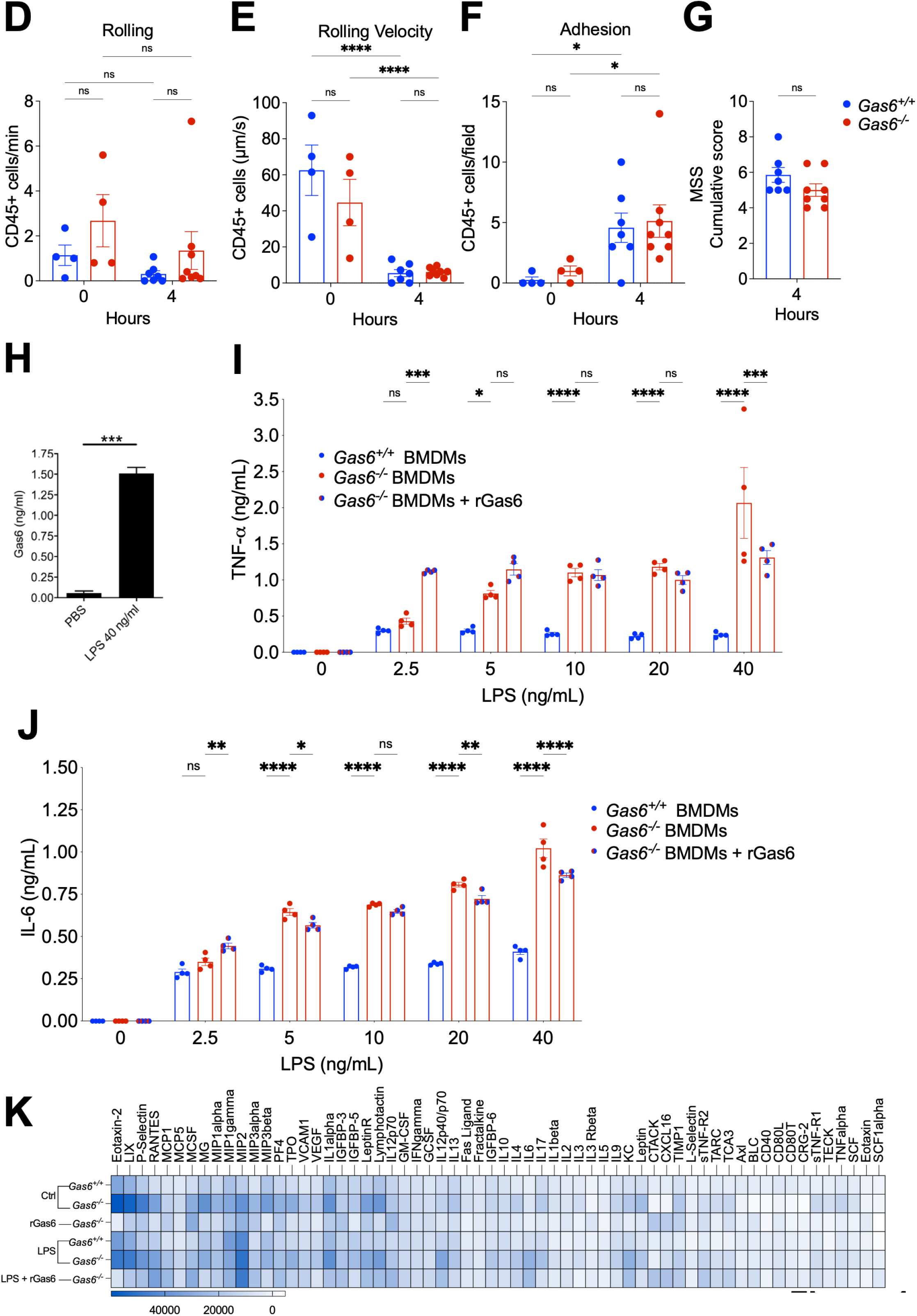
Monocytes/macrophages lacking *Gas6* are activated and overproduce cytokines during endotoxemia. (**A**), Total peripheral blood white blood cell count (WBC), lymphocyte count and monocyte count were assessed in *Gas6^+/+^* and *Gas6^-/-^* mice 24 hours after LPS 50 mg/kg. (**B**), Unsupervised analysis of mass cytometry data of peripheral blood WBC at baseline (controls) and at 1.5 and 4 hours after LPS challenge (10 mg/kg) in *Gas6^+/+^* and *Gas6^-/-^* mice (software: t-SNE). (**C)**, Median marker expression of n = 5-6 mice and absolute cell counts from percentage of parent cell. (**D-F**), Rolling, rolling velocity and adhesion of CD45+ leukocytes in mesenteric post-capillary venules in *Gas6^+/+^* and *Gas6^-/-^* mice at baseline and at 4 hours following LPS challenge (50 mg/kg). (**G**), MSS scores for the intravital microscopy experiment (**H**), Secreted GAS6 by bone marrow-derived macrophages (BMDMs) after 24 hours stimulation with 40 ng/ml LPS. (**I**), TNF-α released by *Gas6^-/-^* bone marrow-derived macrophages (BMDMs) and *Gas6^+/+^* BMDMs after 5-40 ng/ml LPS stimulation. Addition of recombinant murine Gas6 (rGas6, 400 ng/ml) to *Gas6^-/-^* BMDMs conditioned media was able to reduce TNF-α release from *Gas6^-/-^* BMDMs. (**J**), IL-6 secretion by *Gas6^-/-^* BMDMs and *Gas6^+/+^* BMDMs after LPS treatment; addition of rGas6 (400 ng/ml) to the cell culture media reduced IL6 released by *Gas6^-/-^* BMDMs. (**K**), Cytokines antibody array was performed in a pool of 4 samples of culture supernatants from LPS-stimulated BMDMs (40 ng/ml) from *Gas6^-/-^* and *Gas6^+/+^* mice. *Gas6^-/-^* BMDMs media treated with 400 ng/ml rGas6 was also tested. The relative expression of the 62 cytokines has been assessed in arbitrary units (AU) and show in a heat map. Details about the cytokines are listed in **Table 1**. Details about the antibodies are listed in **Table 2**. ns, non significant, \**P* < 0.05, \*\**P* < 0.01, \*\*\**P* < 0.001,\*\*\*\**P* < 0.0001

Overall, the absence of Gas6 did not markedly affect baseline leukocyte composition but altered the early immune dynamics following LPS challenge. Gas6 deficiency was associated with delayed recovery of circulating monocytes and an earlier neutrophil expansion, suggesting a shift toward a more rapid innate inflammatory response. In addition, early increases in T helper and cytotoxic T cells observed in *Gas6^+/+^* mice were attenuated in *Gas6^−/−^* mice, indicating that Gas6 may contribute to the early coordination of adaptive immune cell responses during systemic inflammation.

We next examined the expression intensity of selected monocyte markers. At steady state, Ly6C expression was higher in *Gas6^+/+^* monocytes than in *Gas6^−/−^* monocytes. Following LPS challenge, Ly6C expression rapidly decreased in *Gas6^+/+^* monocytes, reaching levels comparable to those observed in *Gas6^−/−^* monocytes at steady state. At 1.5 hours, Ly6C expression was already lower in *Gas6^+/+^* than in *Gas6^−/−^* monocytes. Reduced expression persisted in both genotypes at 4 hours, although *Gas6^−/−^* monocytes retained slightly higher Ly6C levels than *Gas6^+/+^* monocytes.

These findings suggest that, at steady state, *Gas6^+/+^* mice harbor a higher proportion of Ly6Chigh inflammatory monocytes compared with *Gas6^−/−^* mice, whereas LPS stimulation rapidly normalizes Ly6C expression between genotypes.

Consistently, expression of CD192, which was higher in *Gas6^+/+^* than in *Gas6^−/−^* monocytes at steady state, decreased markedly in both genotypes following LPS challenge and was nearly absent at 1.5 and 4 hours. High CD192 expression is characteristic of inflammatory monocytes and reflects their capacity to migrate from the circulation to inflamed tissues.

Consistent with the changes observed for Ly6C and CD192, the chemokine receptor CX3CR1 was expressed on monocytes from both *Gas6^+/+^* and *Gas6^−/−^* mice at steady state, with slightly higher expression in *Gas6^+/+^* than in *Gas6^−/−^* monocytes. Following LPS challenge, however, CX3CR1 expression was no longer detectable at 1.5 and 4 hours. This might be explained by the fact that CX3CR1 is typically highly expressed on patrolling monocytes (i.e. Ly6Clow monocytes).

Monocytes from both *Gas6^+/+^* and *Gas6^−/−^* mice expressed CD11b at baseline and at 1.5 and 4 hours after LPS administration. CD11b expression increased over time following LPS challenge in both genotypes, reaching its highest levels at 4 hours. Notably, the upregulation of CD11b was more pronounced in *Gas6^+/+^* than in *Gas6^−/−^* monocytes. This increase is consistent with the activated state of circulating monocytes during endotoxemia, as CD11b is a key marker of leukocyte activation and adhesion.

Similarly, neutrophils from both genotypes expressed CD11b and Ly6C at steady state, and the expression of these markers increased during the LPS challenge. At 4 hours, both markers were more strongly expressed on neutrophils from *Gas6^+/+^* mice than on those from *Gas6^−/−^* mice.

In contrast, MHC class II was highly expressed on B cells from both *Gas6^+/+^* and *Gas6^−/−^* mice at steady state and throughout the LPS challenge. However, at 4 hours after LPS administration, MHC class II expression was significantly higher in *Gas6^+/+^* B cells compared with *Gas6^−/−^* B cells.

Together, these data suggest that Gas6 influences baseline leukocyte phenotypes and the early kinetics of the systemic response to LPS. At steady state, circulating monocytes from *Gas6^+/+^* mice exhibited higher Ly6C and CD192 expression than those from *Gas6^−/−^* mice, consistent with a more inflammatory monocyte phenotype. Following LPS challenge, however, Ly6C and CD192 expression rapidly declined in monocytes from both genotypes, with CD192 becoming nearly undetectable by 1.5 and 4 hours. In parallel, CX3CR1 expression disappeared on circulating monocytes in both genotypes, whereas CD11b expression increased over time, reaching highest levels at 4 hours and showing a more pronounced upregulation in *Gas6^+/+^* monocytes.

Beyond monocytes, Gas6 deficiency was associated with an earlier rise in circulating neutrophils, and CD11b and Ly6C expression on neutrophils increased during LPS challenge in both genotypes, with higher expression in *Gas6^+/+^* neutrophils at 4 hours. Finally, B cells maintained high MHC class II expression throughout the time course, but *Gas6^+/+^* B cells displayed significantly higher MHCII levels than *Gas6^−/−^* B cells at 4 hours. Collectively, these findings indicate that while LPS rapidly induces a convergent inflammatory remodeling of circulating monocytes, genotype-dependent differences persist in the magnitude and timing of myeloid activation and in antigen-presentation marker expression on B cells during early endotoxemia.

Although leukocyte rolling tended to be higher in *Gas6^−/−^* mice 4 hours after LPS administration (50 mg/kg), this difference did not reach statistical significance (**Fig. 4D**). No differences were observed in leukocyte rolling velocity or adhesion between genotypes (**Fig. 4E–F**). Importantly, MSS scores were comparable in both genotypes (**Fig. 4G**).

Gas6 and TAM receptors are expressed by macrophages, platelets and endothelial cells but not by circulating lymphocytes and neutrophils^68,69^. Therefore, we focused our next investigations on macrophages as they may represent the main source of circulating Gas6 and cytokines during sepsis^36,70^.

BMDMs released Gas6 upon stimulation with LPS (**Fig. 4H**). We next assessed the functional consequences of *Gas6* deficiency by measuring TNF-α release in *Gas6^+/+^* and *Gas6^-/-^* BMDMs stimulated with LPS. *Gas6^-/-^* BMDMs produced significantly higher levels of TNF-α than *Gas6^+/+^* BMDMs in a dose-dependent manner (**Fig. 4I**). Addition of rGas6 (400 ng/mL) to *Gas6^-/-^* BMDMs reduced TNF-α release from *Gas6^-/-^* BMDMs, thereby restoring a *Gas6^+/+^* phenotype. A similar restoration of the *Gas6^+/+^* phenotype was observed in time-course experiments. Likewise, IL-6 secretion was significantly increased in *Gas6^-/-^* BMDMs compared to *Gas6^+/+^* BMDMs and its and increased progressively with LPS dose (**Fig. 4J**). Increased IL-6 levels in *Gas6^-/-^* BMDMs were dampened by adding rGas6 (400 ng/mL) to the cell culture medium (**Fig. 4J**). To further expand these findings, we performed a cytokine array quantifying 62 inflammatory and immunomodulatory mediators in culture supernatants after 7 hours of LPS stimulation. *Gas6^-/-^* BMDMs released higher levels of multipled cytokines compared with *Gas6^+/+^* BMDMs (**Fig. 4K**). Addition of rGas6 normalized cytokine production, restoring levels comparable to those observed in *Gas6^+/+^* macrophages.

Together, these results indicate that macrophages produce Gas6 in response to inflammatory stimulation and that Gas6 acts as an autocrine regulatory signal limiting excessive cytokine release. In the absence of Gas6, macrophages become hyperresponsive to LPS stimulation and produce higher levels of inflammatory mediators. Restoration of Gas6 signaling with recombinant Gas6 largely reverses this phenotype, supporting a model in which Gas6–TAM signaling acts as an endogenous negative feedback mechanism controlling macrophage activation and inflammatory cytokine production during endotoxemia.

### Gas6 immunomodulatory function acted through SOCS1/3 pathway in monocytes

Signal transduction mediated by Gas6 in other cell types than dendritic cells is known to involve the activation of PI3K and its downstream target, Akt^71^. In addition, Mer regulates PI3K and NF-κB activation in dendritic cells^72^. Since it has been shown that the PI3K/Akt signaling pathway negatively regulates LPS-induced acute inflammation^73^, we assessed whether the absence of *Gas6* affects Akt activation in BMDMs. Moreover, downstream targets of the PI3K/Akt pathway and the TLR4 pathway include the MAPK pathways (p38, ERK1/2, JNK)^74^. Thus, we stimulated BMDMs with LPS (40 ng/mL) and assessed the phosphorylation of Akt, p38, ERK1/2 and JNK, as well as the total form of each protein (**Fig. 5A**). Consequently, we did not notice a modification of the phosphorylation state of IκB-α between *Gas6^+/+^* and *Gas6^-/-^* BMDMs (**Fig. 5B**).

**FIGURE 5:**
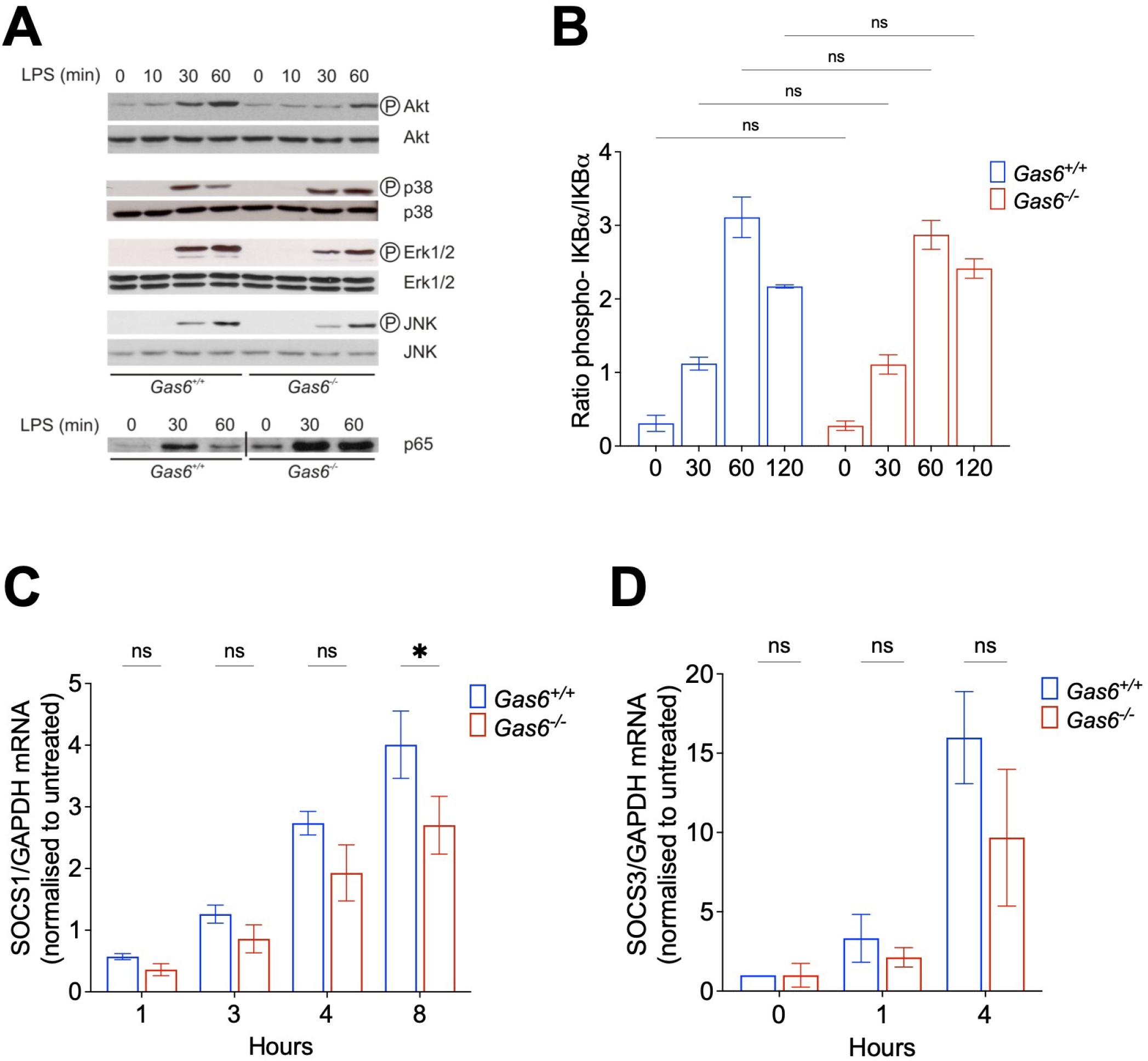
SOCS1 pathway is dampend in *Gas6^-/-^* macrophages. (**A**), Time course of Akt, p38, ERK1/2 and JNK phosphorylation in response to 40 ng/ml lipopolysaccharide (LPS) in bone marrow-derived macrophages (BMDMs). Representative blot from 3 independent experiments. (**B**), Kinetics of IκB phosphorylation after LPS (40 ng/ml) stimulation in BMDMs (3 independent experiments). (**C**), Relative mRNA expression of suppressor of cytokine signaling (SOCS)1 quantified by qPCR using glyceraldehyde 3-phosphate dehydrogenase (GAPDH) as housekeeping gene and normalized to non-stimulated cells (0 hour). BMDMs were incubated for 1 to 8 hours with 40 ng/ml of LPS. Tukey’s multiple comparisons test. (**D**), Relative mRNA expression of SOCS3 quantified by qPCR using GAPDH as housekeeping gene and normalized with baseline expression. BMDMs were incubated for 1 to 4 hours with 40 ng/ml of LPS. \**P* < 0.05, \*\**P* < 0.01

Gas6 induces the cytokine and TLR suppressors SOCS1 and SOCS3 in dendritic cells, as shown by an increased mRNA expression of SOCS1/3 in cell lysates^52^ and in the synovia of mice in a collagen-Induced arthritis model^75^. Quantitative RT-PCR of BMDMs stimulated with LPS at different time points was performed. In absence of Gas6, mRNA expression of SOCS1 was reduced (**Fig. 5C**). A similar pattern was also observed for SOCS3 although only as a trend (**Fig. 5D**). The same pattern has been described in^75^. Thus, the lack of Gas6 weakened the regulation of the TLR signaling pathway in BMDMs by dampening at least SOCS1 activation.

### Sepsis-induced coagulopathy is associated with elevated Gas6 and sMer levels

Sepsis-induced coagulopathy (SIC) is a frequent complication in septic patients and is characterized by systemic inflammation accompanied by activation of the coagulation cascade. Thrombin generation plays a central role in coagulation activation during sepsis and exerts multiple cellular effects, thereby contributing substantially to sepsis pathophysiology ^76,77^. However, conventional coagulation assays, such as prothrombin time (PT) and activated partial thromboplastin time (aPTT), are insufficient to reliably assess thrombotic or hemorrhagic phenotypes. Measurement of thrombin-generating capacity may therefore provide a more comprehensive assessment of coagulation status than routine coagulation tests ^78^. Because blood cells critically contribute to coagulation and influence thrombin generation (TG) testing, whole blood TG offers a more physiological evaluation than plasma-based assays, as it incorporates the contributions of red blood cells, leukocytes, and platelets ^79,80^.

We first calculated the SIC score in 20 patients within 24 hours of admission for septic shock and in healthy volunteers (n = 6). Patients with SIC (score ≥ 4) exhibited significantly higher plasma Gas6 levels than those with a score < 4 (**Fig. 6A**). Furthermore, SIC scores positively correlated with plasma Gas6 and sMer levels, but not with sAxl or sTyro3 (**Fig. 6B**).

**FIGURE 6:**
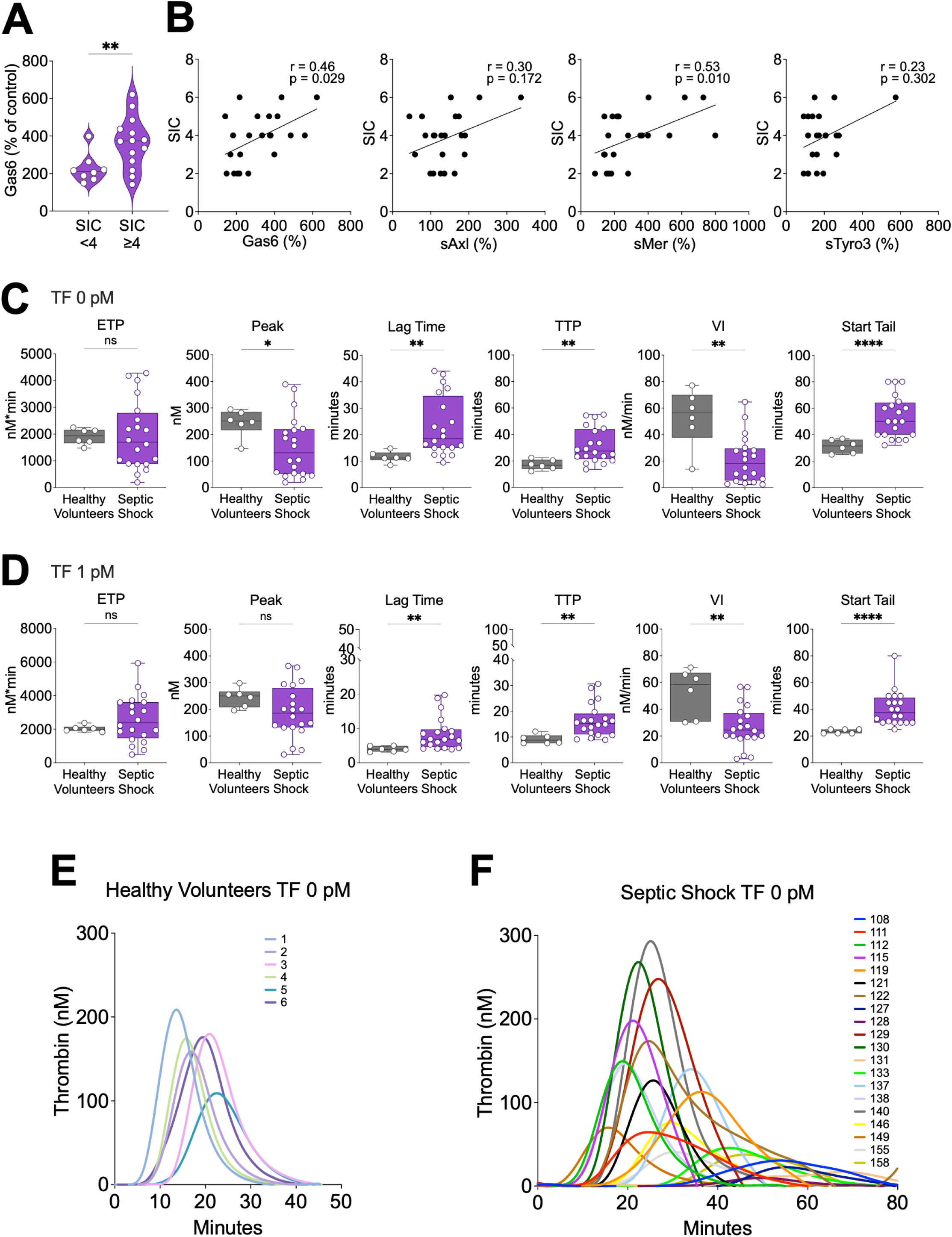
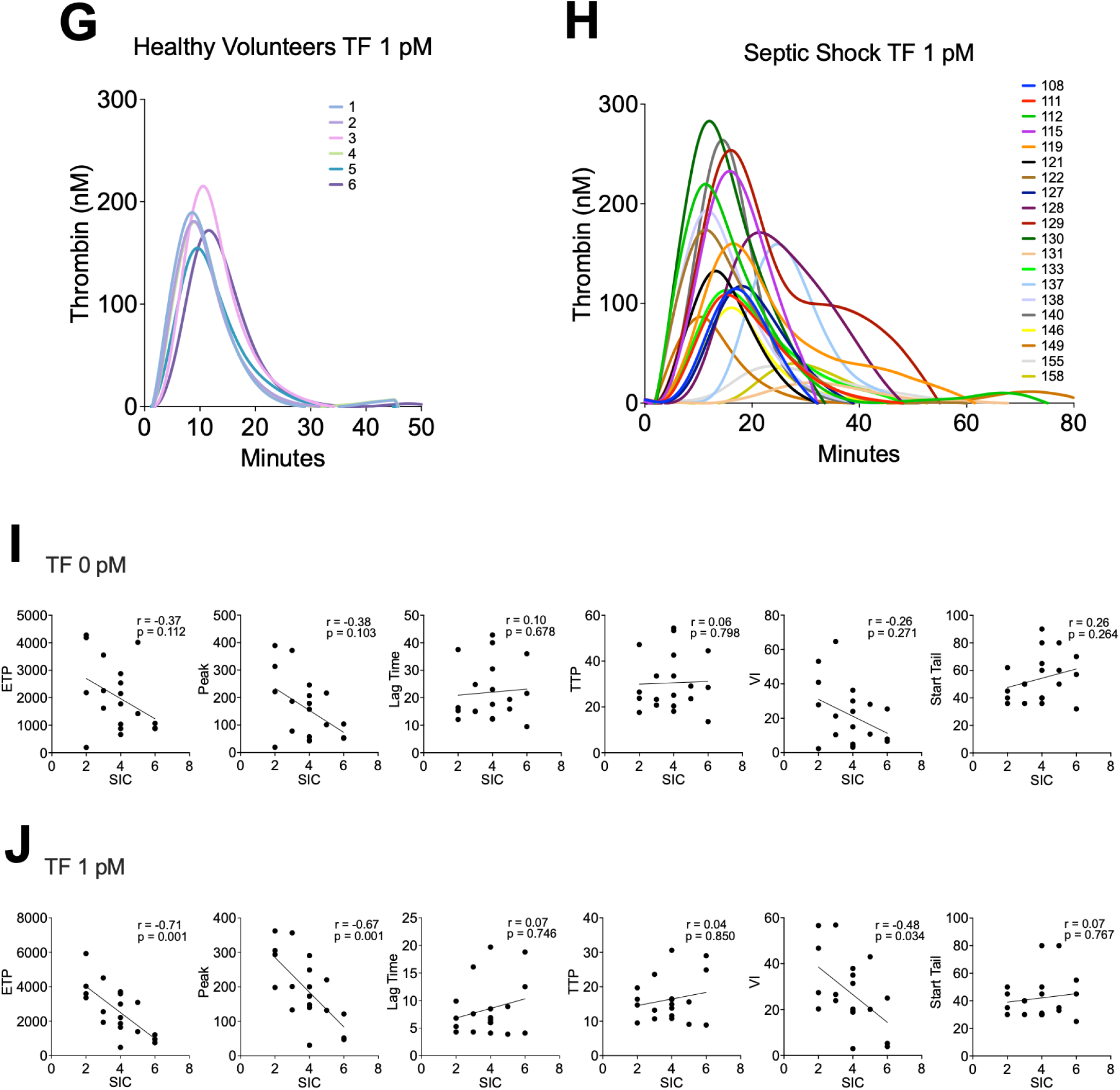
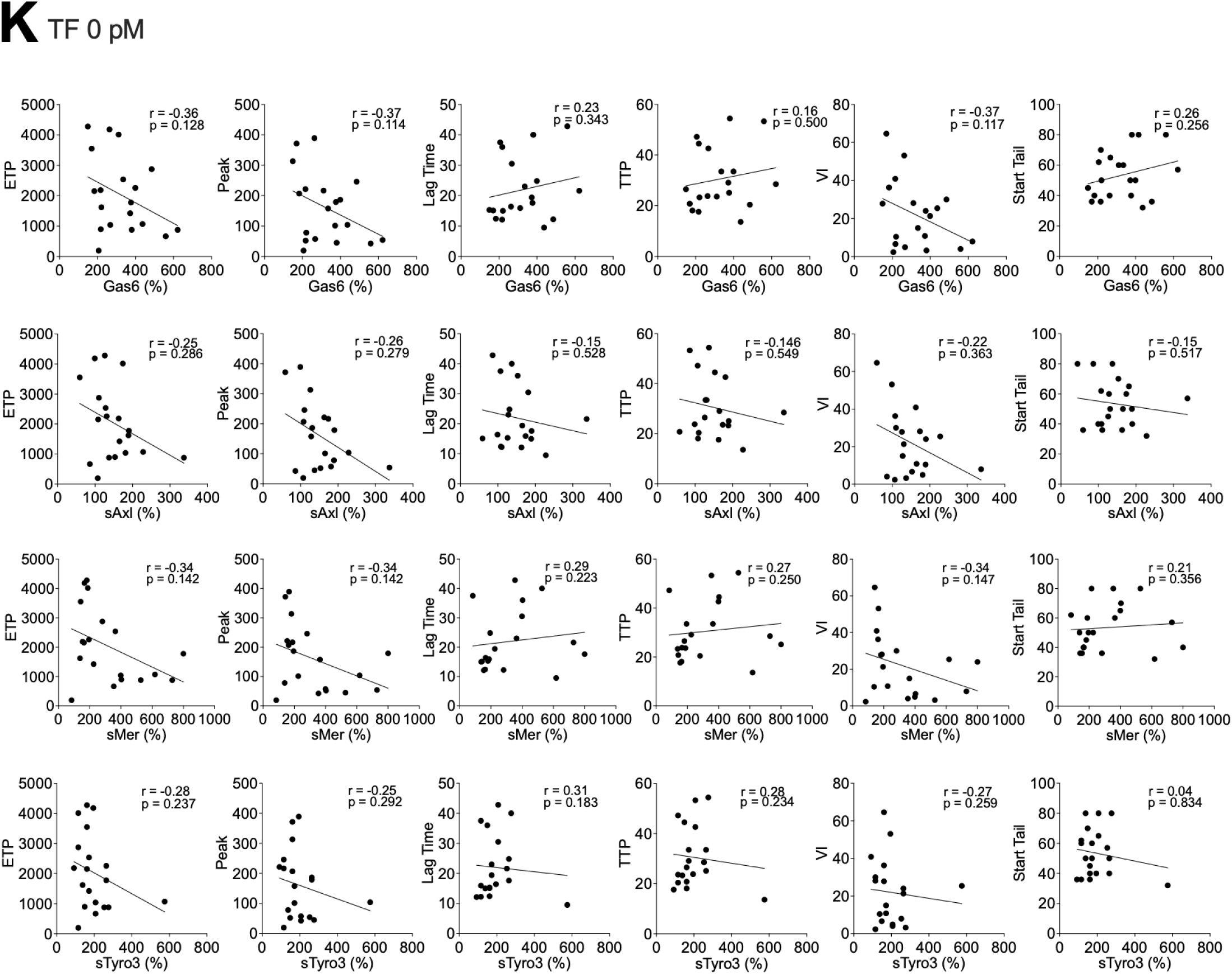
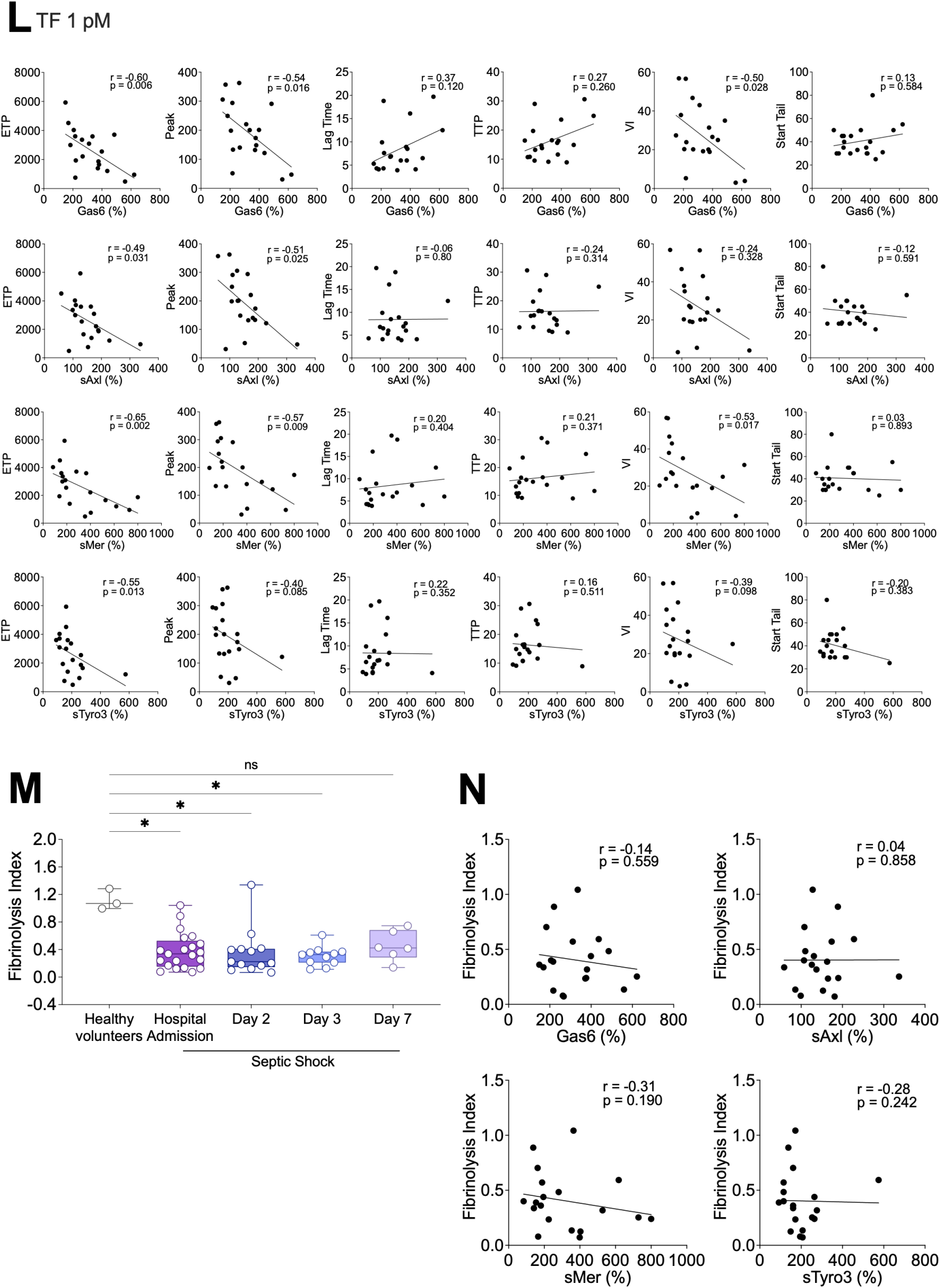
Sepsis-induced coagulopathy is associated with elevated Gas6 and sMer levels. (**A**), Gas6 plasma levels in patients with septic shock and either a sepsis-induced coagulopathy score (SIC) <4 or ≥4. (**B**), Spearman’s correlation and linear regression between SIC score and respectively Gas6, sAxl, sMer and sTyro3 plasma levels. (**C-H**), Thrombin generation in whole blood in healthy volunteers and in patients with septic shock, tissue factor (TF) 0 pM (**C**) or 1 pM (**D**), with mean thrombograms (**E-H**). (**I-J**), Spearman’s correlation and linear regression between the different thrombin generation parameters (**I**: TF 0 pM and **J**: TF 1 pM) and SIC score. (**K-L**), Spearman’s correlation and linear regression between thrombin generation parameters and respectively Gas6, sAxl, sMer and sTyro3 plasma levels (**K**: TF 0 pM and **L**: TF 1 pM). (**M**), Fibrinolysis index measured in healthy volunteers and patients with septic shock at admission and Day 2, Day 3 and Day 4. (N), Spearman’s correlation and linear regression between fibrinolysis index and Gas6, sAxl, sMer and sTyro3, respectively. \**P* < 0.05, \*\**P* < 0.01, \*\*\**P* < 0.001,\*\*\*\**P* < 0.0001

Whole blood TG was then assessed in septic shock patients and healthy volunteers, both in the absence (**Fig. 6C**) and presence of tissue factor (TF, 1 pM) (**Fig. 6D**). In the absence of TF, endogenous thrombin potential (ETP) did not differ significantly between groups. However, thrombin peak and velocity index were reduced in septic shock patients, whereas lag time and time to peak were prolonged. In the presence of TF, lag time and time to peak remained prolonged, and velocity index remained reduced in septic shock patients compared with controls. These findings indicate that thrombin generation is delayed and proceeds more slowly in septic shock. This pattern is consistent with previous plasma-based TG measurements during infection ^81^. Importantly, start tail was markedly prolonged in septic shock patients compared with healthy volunteers, both without and with TF triggering (**Fig. 6C–D**), suggesting delayed thrombin termination and further supporting the presence of SIC^82^. Thrombograms are shown in **Fig. 6E–H**. Greater inter-individual variability was observed among septic shock patients than among healthy controls, particularly in the absence of TF. Thrombograms from septic patients were markedly larger, reflecting the heterogeneity of coagulation responses during septic shock.

We next examined correlations between TG parameters (ETP, thrombin peak, lag time, time to peak, velocity index, and start tail) and SIC scores (**Fig. 6I-J**). Without TF triggering, no significant correlations were observed (**Fig. 6I**). In contrast, when TG was triggered with 1 pM TF, ETP, thrombin peak, and velocity index were inversely correlated with SIC score, whereas lag time, time to peak, and start tail showed no significant associations (**Fig. 6J**).

We further analyzed correlations between TG parameters and circulating levels of Gas6, sAxl, sMer, and sTyro3 (**Fig. 6K-L**). In the absence of TF, no significant correlations were detected (**Fig. 6K**). However, with 1 pM TF triggering, ETP inversely correlated with Gas6, sAxl, sMer, and sTyro3 levels. Thrombin peak inversely correlated with Gas6, sAxl, and sMer, while velocity index inversely correlated with Gas6 and sMer (**Fig. 6L**).

In septic shock, alongside dysregulated thrombin generation and impaired termination, defective fibrinolysis restricts fibrin degradation and promotes persistent microvascular thrombosis—the pathological hallmark of sepsis-associated disseminated intravascular coagulation ^83^. The fibrinolysis index was significantly reduced in septic patients compared with healthy volunteers (**Fig. 6M**), indicating impaired fibrinolytic capacity. This finding is consistent with the well-established hypofibrinolytic state in sepsis, often referred to as “fibrinolytic shutdown,” largely driven by elevated plasminogen activator inhibitor-1 levels ^84^. However, unlike TG parameters, the fibrinolysis index did not correlate with circulating Gas6 or soluble TAM receptor levels (**Fig. 6L**).

Taken together, these data demonstrate that plasma Gas6 levels are significantly higher in septic shock patients with SIC than in those without SIC, and that SIC scores positively correlate with circulating Gas6 and sMer, but not with sAxl or sTyro3. Septic shock was characterized by delayed and reduced thrombin generation, together with prolonged thrombin termination, indicating a dysregulated and heterogeneous coagulation response consistent with sepsis-induced coagulopathy. Notably, TF-triggered thrombin generation parameters (ETP, thrombin peak, and velocity index) were inversely associated with SIC score as well as with circulating Gas6 and soluble TAM receptor levels, particularly sMer, suggesting a link between TAM pathway activation and the severity of coagulation impairment. Although hypofibrinolysis was evident in our septic cohort, fibrinolytic capacity was not associated with Gas6 or soluble TAM receptor concentrations, indicating that the relationship between the Gas6/TAM axis and SIC primarily involves thrombin generation rather than fibrinolytic shutdown. These findings extend our previous observation of elevated plasma Gas6 levels in septic patients and further demonstrate that both Gas6 and sMer correlate with SIC severity. Moreover, they are in line with published studies showing that delayed and reduced thrombin generation predicts adverse outcomes in sepsis ^81^, thereby supporting the clinical relevance of impaired thrombin dynamics in this setting.

### Limited impact of Gas6 on coagulation activation during endotoxemia in mice

Gas6 is generally considered a procoagulant protein^55,85-88^. We previously reported that elevated plasma Gas6 levels predict venous thromboembolism recurrence, major bleeding, and mortality in elderly patients^88^. In the present study, however, we observed that high Gas6 levels in septic patients correlated with reduced thrombin generation. This apparent paradox suggests that the relationship between Gas6 and coagulation in sepsis may differ from its role in other clinical contexts. Although elevated Gas6 levels in sepsis are associated with adverse outcomes, this association does not establish a direct mechanistic link between elevated Gas6 in plasma and coagulation impairment during sepsis.

To address this question, we investigated the coagulation response to systemic inflammation *in vivo* using *Gas6^+/+^* and *Gas6^−/−^* mice challenged with LPS, as we have previously shown that plasma Gas6 levels increase during endotoxemia in mice, similarly to what is observed in humans.

We first assessed classical parameters associated with DIC. Platelet counts, prothrombin time, fibrinogen levels and thrombin–antithrombin (TAT) complexes were measured in *Gas6^+/+^* and *Gas6^−/−^* mice 24 hours after injection of LPS (50 mg/kg). These parameters were comparable between the two genotypes, except for platelet counts, which were significantly higher in *Gas6^−/−^* than in *Gas6^+/+^* mice, suggesting reduced platelet consumption in *Gas6^−/−^* mice following LPS challenge (**Fig. 7A**).

**FIGURE 7:**
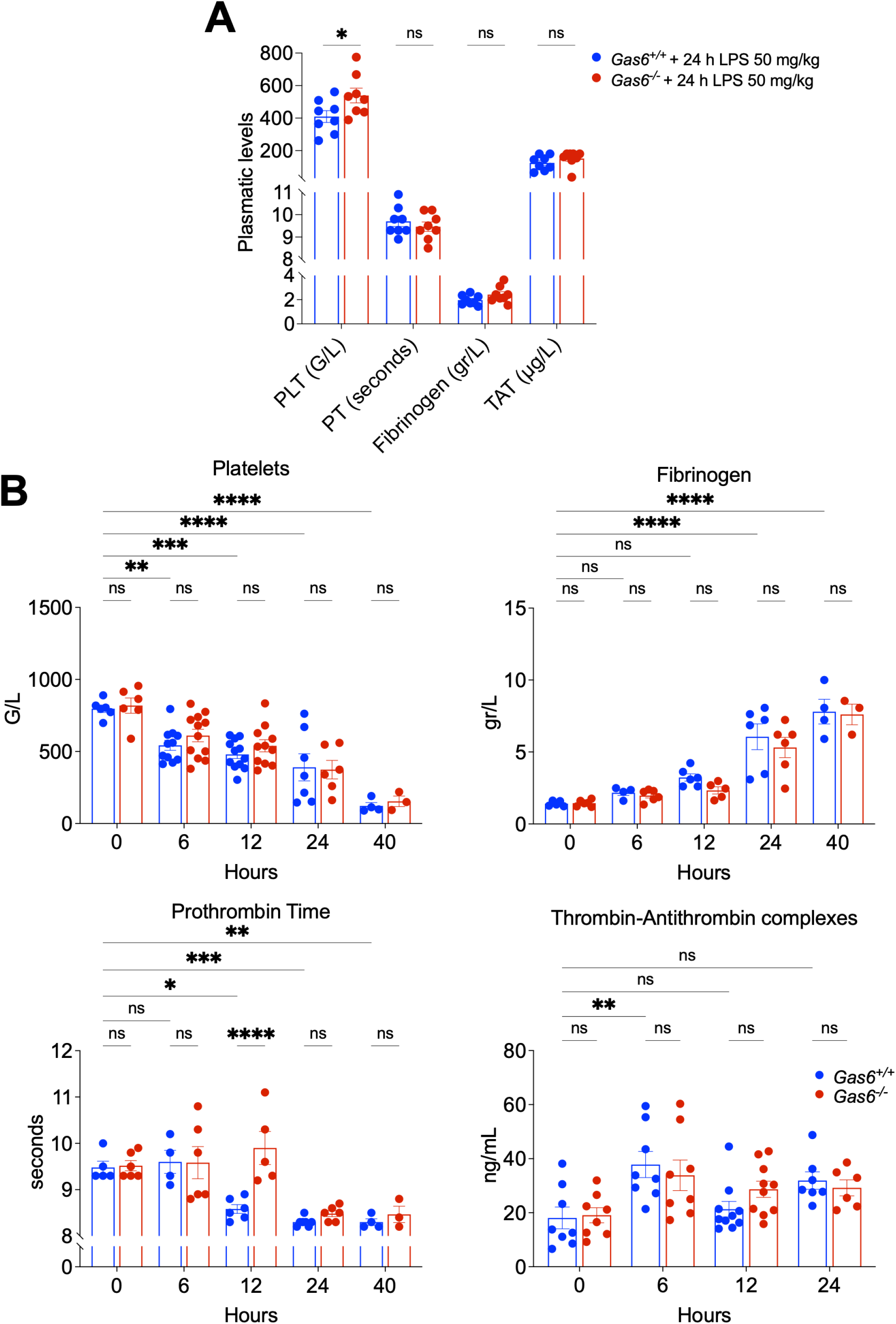
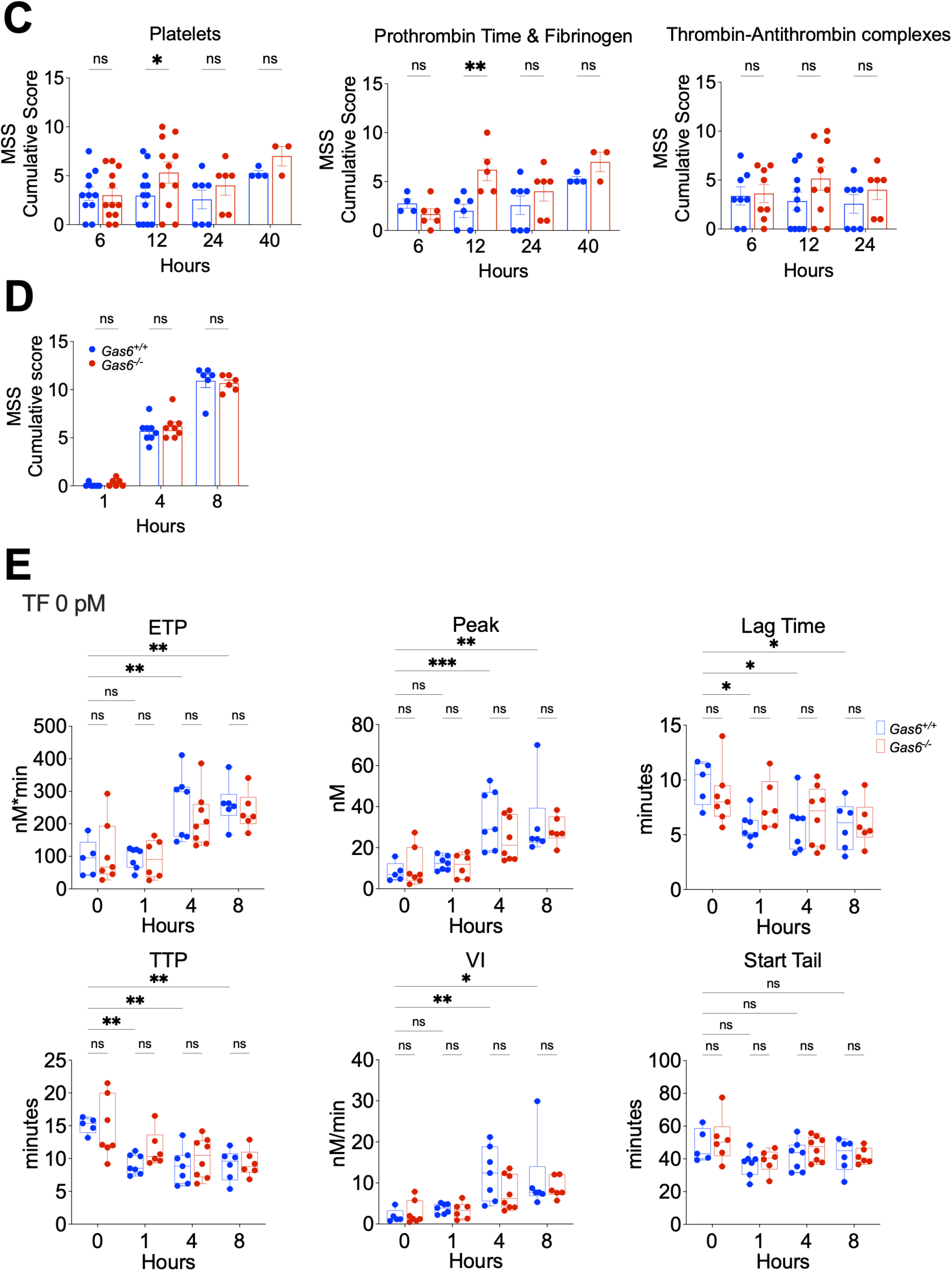
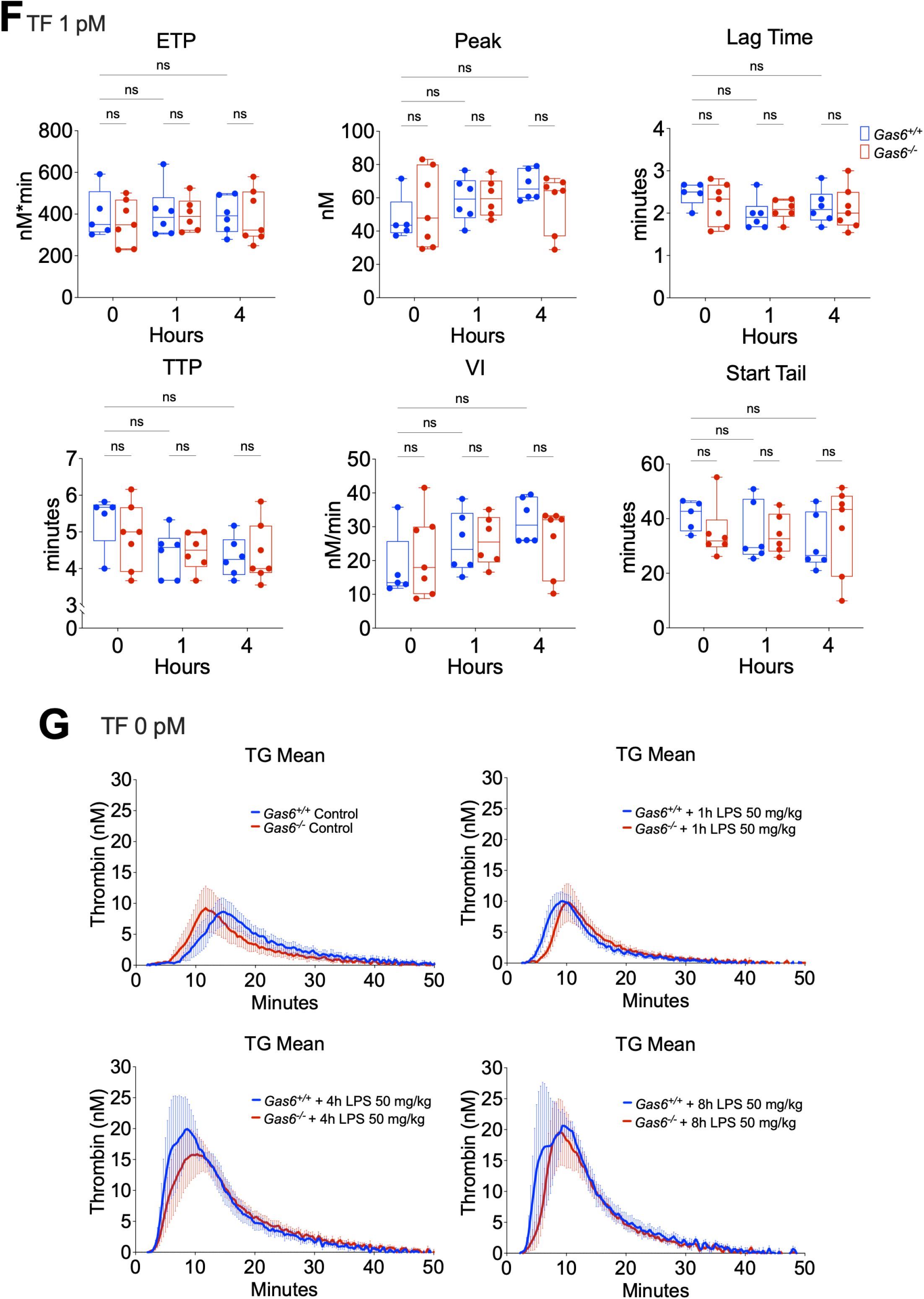
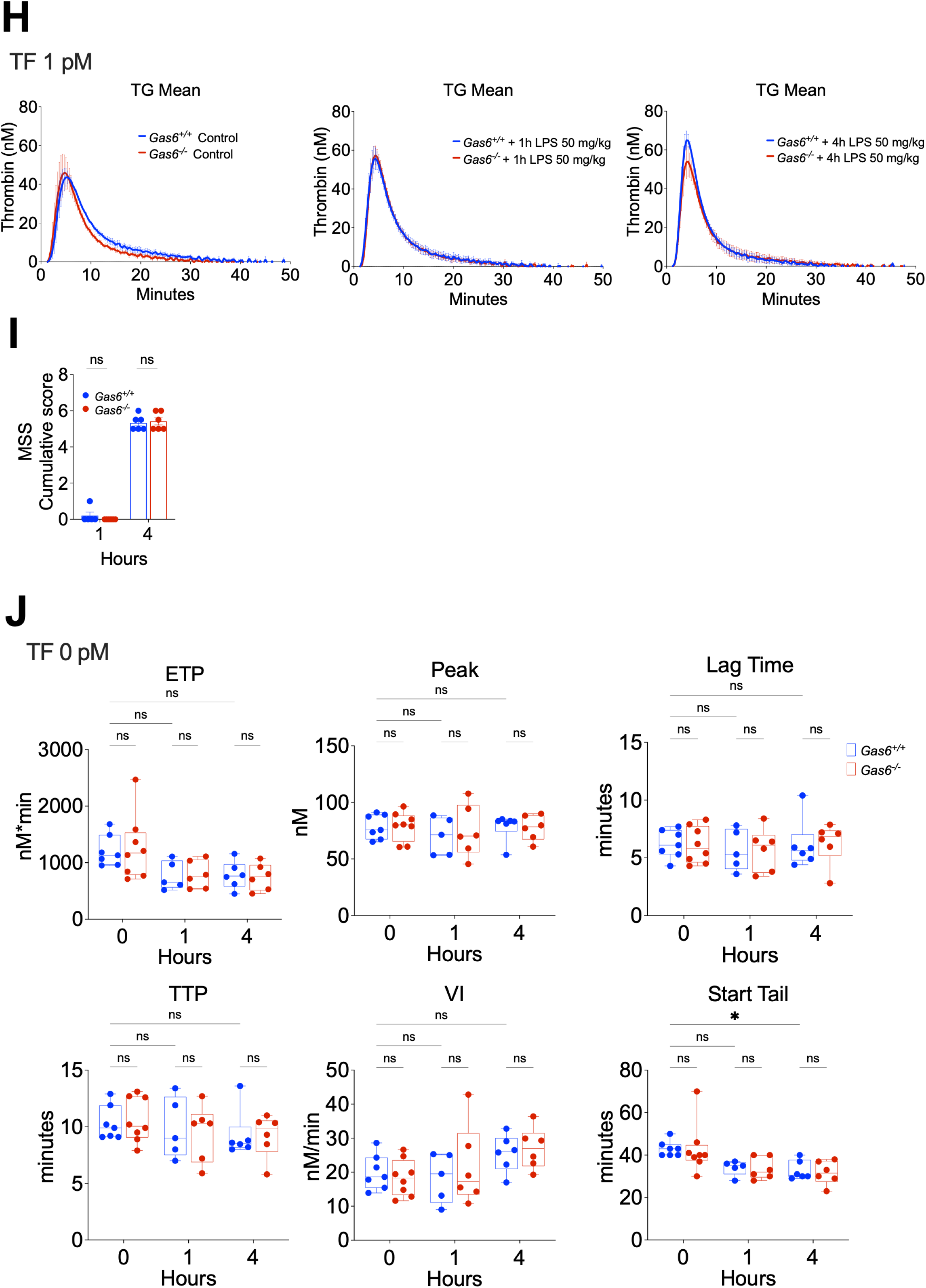
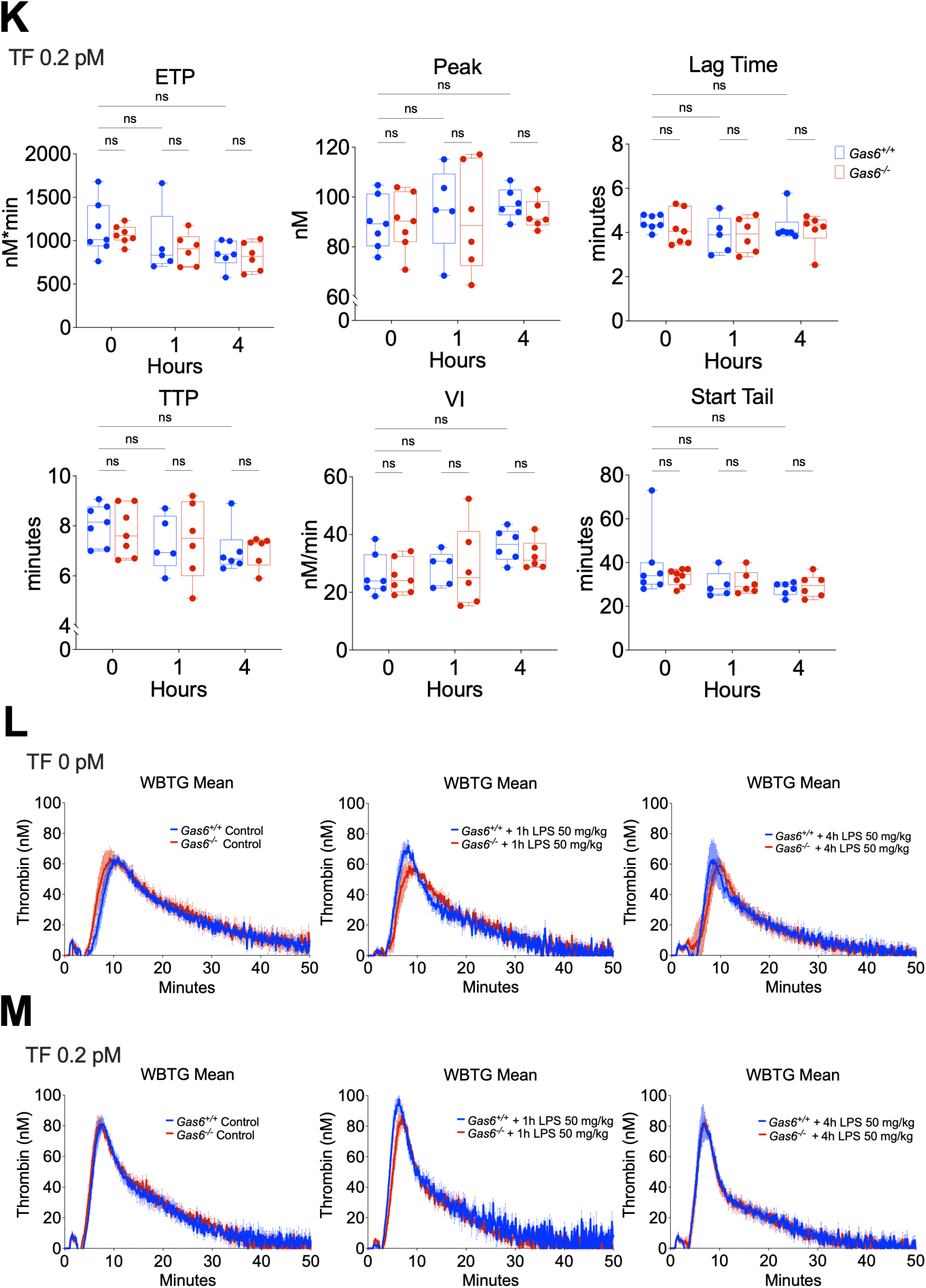
Limited impact of Gas6 on coagulation activation during endotoxemia in mice. (**A**), Disseminated intravascular coagulation parameters (DIC): platelet count (PLT), prothrombin time (PT), fibrinogen, thrombin-antithrombin complexes measured in plasma of *Gas6^+/+^* and *Gas6^-/-^* mice 24 hours after LPS 50 mg/kg. (**B-C**), Disseminated intravascular coagulation parameters (DIC): platelet count (PLT), prothrombin time (PT), fibrinogen, thrombin-antithrombin complexes measured in plasma of *Gas6^+/+^* and *Gas6^-/-^* mice at different time points after LPS 10 mg/ml under heating conditions (**B**) and MSS scores collected at the different timepoints (**C**). (**D-H**), Murine sepsis score (MSS) scores (**D**), thrombin generation in whole blood, tissue factor (TF) 0 pM measured at different time points after LPS 50 mg/kg (**E**), thrombin generation in whole blood, tissue factor (TF) 1 pM measured at different time points after LPS 50 mg/kg (**F**), Mean thrombograms (**G-H**). (**I-M**), MSS scores (**I**), thrombin generation in platelet-free plasma, tissue factor (TF) 0 pM measured at different time points after LPS 50 mg/kg (**J**), thrombin generation in platelet-free plasma, tissue factor (TF) 1 pM measured at different time points after LPS 50 mg/kg (**K**), Mean thrombograms (**L-M**). \**P* < 0.05, \*\**P* < 0.01, \*\*\**P* < 0.001,\*\*\*\**P* < 0.0001

In a milder endotoxemia model (LPS 10 mg/kg) (**Fig. 7B**), platelet counts decreased in both genotypes throughout the experiment (up to 40 hours), but no differences between genotypes were observed. Prothrombin time transiently shortened at 12 hours in *Gas6^+/+^* mice but not in *Gas6^−/−^* mice, in which it remained similar to baseline. However, prothrombin time was comparable between genotypes at later time points (24 and 40 hours). Fibrinogen levels increased at 24 and 40 hours in both genotypes, consistent with an acute-phase response to systemic inflammation. Thrombin–antithrombin complexes increased significantly at 6 hours in *Gas6^+/+^* mice but not in *Gas6^−/−^* mice. Importantly, mouse severity score (MSS) values were comparable between genotypes throughout the experiment, except at 12 hours, where the score was slightly higher in *Gas6^-/-^* than in *Gas6^+/+^* mice (**Fig. 7C**). Overall, these observations suggest a trend toward reduced coagulation activation in *Gas6^−/−^* mice compared with Gas6+/+ mice. Nevertheless, neither genotype developed overt DIC during endotoxemia.

To further characterize coagulation activation during LPS challenge, thrombin generation (TG) assays were performed in platelet-poor plasma (PPP) from *Gas6^+/+^* and *Gas6^−/−^* mice, either without tissue factor (TF) trigger or with TF (1 pM) with follow-up of MSS scores (**Fig. 7D-H**). In the absence of TF (**Fig. 7E**), several TG parameters changed during endotoxemia in both genotypes, consistent with systemic coagulation activation and the increase in MSS scores (**Fig. 7E**). These changes included increased endogenous thrombin potential (ETP) and thrombin peak, a modest reduction in lag time and time to peak, and an increase in velocity index, whereas start tail remained unchanged. Importantly, all TG parameters were comparable between *Gas6^+/+^* and *Gas6^−/−^* mice throughout the 8-hour observation period. When TG was triggered with TF (1 pM) (**Fig. 7F**), no significant changes in TG parameters were observed during the 4-hour observation period, and results remained comparable between the two genotypes, despite increase MSS scores upon LPS challenge (**Fig. 7D**). Representative thrombograms for both conditions (with and without TF) are shown in **Fig. 7G-H**. Similarly, thrombin generation measured in whole blood revealed no significant differences between genotypes or time points (**Fig. 7I-K**), despite a significant increase in MSS scores at 4 hours in both groups (**Fig. 7I**). Representative thrombograms obtained under these conditions are shown in **Fig. 7L-M**.

Taken together, these results indicate that although endotoxemia induces signs of coagulation activation at later time points, neither *Gas6^+/+^* nor *Gas6^−/−^* mice develop disseminated intravascular coagulation during LPS challenge. While Gas6 deficiency is associated with a modest reduction in markers of coagulation activation, the overall coagulation response to endotoxemia remains largely comparable between genotypes.

## Discussion

In this study, we investigated the role of Gas6–TAM signaling during systemic inflammation and sepsis using complementary clinical observations in humans and experimental models in mice. Our findings identify Gas6 as an important endogenous regulator of inflammatory responses during endotoxemia and infection. Plasma Gas6 levels increased rapidly during experimental endotoxemia in mice and in patients with sepsis, correlated with disease severity, and were associated with adverse clinical outcomes. Despite this pronounced increase, circulating Gas6 remained closely balanced with sAxl, suggesting that a large proportion of the ligand is sequestered within soluble Gas6–sAxl complexes. This buffering mechanism likely limits the amount of free Gas6 available to activate membrane-bound TAM receptors.

Our data further demonstrate that Gas6 influences host responses to endotoxemia and bacterial infection through modulation of innate immune signaling. Functional experiments showed that Gas6 deficiency markedly increased susceptibility to endotoxemia and bacterial infection, supporting a protective role for endogenous Gas6 during systemic inflammation. These findings are consistent with previous studies demonstrating that TAM receptor signaling acts as an important negative regulator of innate immune activation. For example, activation of Mer has been shown to suppress TNF-α production by monocytes and macrophages and to attenuate endotoxic shock in mice^49^. Likewise, macrophages lacking all three TAM receptors (Tyro3, Axl, and Mer) exhibit chronic hyperactivation and excessive production of pro-inflammatory cytokines following endotoxin stimulation^56^. In dendritic cells, TAM signaling has been reported to provide a negative feedback mechanism that limits TLR- and cytokine-driven immune responses^52^.

Clinical observations further support the involvement of the Gas6–TAM axis in sepsis pathophysiology. Early measurement of sMer and Gas6 in patients presenting with sepsis did not predict short-term mortality but was associated with sepsis-related organ dysfunction, particularly acute kidney injury and coagulation abnormalities, suggesting that these biomarkers may reflect early organ damage rather than overall prognosis^42^. In addition, experimental studies have shown that administration of Gas6 in combination with antibiotics reduces organ injury, inflammatory responses, and oxidative stress in murine models of sepsis, indicating that Gas6 supplementation may have therapeutic potential in this context^47^. Based on these observations, we hypothesized that endogenous Gas6 may contribute to protection against the systemic inflammatory response to infection and potentially influence the immune dysregulation observed in severe sepsis.

Consistent with this hypothesis, *Gas6^−/−^* mice exhibited exaggerated systemic inflammation characterized by elevated cytokine levels, reduced arterial blood pressure, and altered leukocyte dynamics, including delayed monocyte recovery and accelerated neutrophil expansion. At the cellular level, macrophages released Gas6 in response to LPS stimulation, and Gas6 acted as an autocrine negative feedback signal limiting inflammatory cytokine production. In the absence of Gas6, macrophages became hyperresponsive to inflammatory stimulation, producing higher levels of TNF-α, IL-6, and other inflammatory mediators. Restoration of Gas6 signaling with recombinant Gas6 normalized cytokine production, supporting a model in which Gas6–TAM signaling functions as an endogenous brake on macrophage activation, at least in part through modulation of TLR-dependent pathways.

Our stoichiometric analysis further supports the concept that the circulating Gas6–sAxl system functions as a dynamic buffering compartment regulating ligand availability during inflammation. Previous work by Ekman et al.^64^ proposed that most circulating Gas6 exists in complex with sAxl, forming a high-affinity ligand–receptor reservoir in plasma. Our measurements in *Gas6^+/+^* mice are consistent with this model, as baseline Gas6 and sAxl concentrations were present at near-equimolar levels, theoretically allowing sequestration of the majority of circulating Gas6 in soluble complexes. Importantly, endotoxemia induced parallel increases in both Gas6 and sAxl, maintaining this near-stoichiometric equilibrium and suggesting that inflammatory responses expand the buffering capacity of the soluble Axl compartment rather than simply increasing free ligand availability. Our measurements in healthy volunteers and patients with septic shock further support the concept that circulating Gas6 is largely bound to soluble Axl. In healthy individuals, Gas6, sAxl, and Gas6–sAxl complexes were present at comparable concentrations, consistent with a near-complete sequestration of circulating Gas6 by sAxl under physiological conditions. Although Gas6 levels increased in septic shock, sAxl remained present in slight excess, indicating that the soluble Axl compartment was not fully saturated and that the amount of free Gas6 available to activate membrane TAM receptors is likely limited even during severe inflammation.

In contrast, administration of recombinant Gas6 exceeded this buffering capacity, resulting in a measurable reduction of circulating sAxl and a several-fold increase in calculated free Gas6 concentrations. These observations indicate that exogenous Gas6 can shift the system from a buffered equilibrium toward a ligand-excess state compatible with productive TAM receptor activation. Interestingly, this mechanism aligns with our observations in human sepsis cohorts, where distinct patient groups exhibited either Gas6-dominant or sAxl-dominant plasma states. This suggests that the balance between ligand and soluble decoy receptor may represent an important determinant of TAM pathway activity during sepsis. Together, these findings support a model in which the Gas6–sAxl equilibrium acts as a regulatory rheostat controlling TAM signaling and raise the possibility that therapeutic strategies aimed at increasing Gas6 availability or interfering with its sequestration by sAxl may restore protective TAM signaling under conditions of severe inflammation.

In contrast to its well-established role in immune regulation, our data do not support a major role for Gas6 as a direct regulator of coagulation during endotoxemia.

In patients with septic shock, circulating Gas6 and soluble Mer concentrations were associated with sepsis-induced coagulopathy scores and impaired thrombin generation. However, this association most likely reflects the role of Gas6 as a marker of disease severity rather than a direct mediator of coagulation activation. Because circulating Gas6 remains largely bound to soluble Axl, the fraction of free ligand available to activate membrane TAM receptors on endothelial cells, platelets, or monocytes is expected to remain limited even under septic conditions. This buffering mechanism provides a plausible explanation for the apparent discrepancy between the association of Gas6 with sepsis severity and the absence of a pronounced procoagulant phenotype in vivo. Elevated Gas6 levels may therefore primarily reflect the magnitude of systemic inflammation and endothelial activation rather than directly driving coagulation abnormalities. Although hypofibrinolysis was evident in our septic cohort, fibrinolytic capacity was not associated with Gas6 or soluble TAM receptor concentrations, suggesting that the relationship between the Gas6–TAM axis and sepsis-induced coagulopathy is more closely linked to thrombin generation than to impaired fibrinolysis.

In our murine model, Gas6 deficiency resulted in only modest differences in markers of coagulation activation, and neither *Gas6^+/+^* nor *Gas6^−/−^* mice developed disseminated intravascular coagulation following LPS challenge. Thrombin generation parameters were largely comparable between genotypes, indicating that the overall coagulation response to endotoxemia is not substantially driven by Gas6 signaling in this experimental setting.

Taken together, our findings position Gas6 as an important endogenous modulator of inflammatory responses during sepsis and highlight the Gas6–TAM axis as a marker of disease severity and immune– vascular activation. Moreover, the dynamic balance between Gas6 and its soluble decoy receptor sAxl appears to regulate ligand bioavailability and TAM receptor signaling during systemic inflammation, suggesting that modulation of this axis may represent a potential therapeutic strategy in sepsis.

## Acknowledgments

The authors thank Rocco Sugamele, Monica Azevedo, Béatrice Ternon, Tecla Dudez and Debora Sanches Rodrigues for technical assistance.

